# Integrative analyses of convergent adaptation in sympatric extremophile fishes

**DOI:** 10.1101/2021.06.28.450104

**Authors:** Ryan Greenway, Rishi De-Kayne, Anthony P. Brown, Henry Camarillo, Cassandra Delich, Kerry L. McGowan, Joel Nelson, Lenin Arias-Rodriguez, Joanna L. Kelley, Michael Tobler

## Abstract

The evolution of independent lineages along replicated environmental gradients frequently results in convergent adaptation, yet the degree to which convergence is present across multiple levels of biological organization is often unclear. Additionally, inherent biases associated with shared ancestry and variation in selective regimes across geographic replicates often pose challenges for confidently identifying patterns of convergence. We investigated a system in which three species of poeciliid fishes sympatrically occur in a toxic spring rich in hydrogen sulfide (H_2_S) and an adjacent nonsulfidic stream to examine patterns of adaptive evolution across levels of biological organization. We found convergence in morphological and physiological traits and genome-wide patterns of gene expression among all three species. In addition, there were shared signatures of selection on genes encoding H_2_S toxicity targets in the mitochondrial genomes of each species. However, analyses of nuclear genomes revealed neither evidence for substantial genomic islands of divergence around genes involved in H_2_S toxicity and detoxification nor substantial congruence of strongly differentiated regions across population pairs. These non-convergent, heterogenous patterns of genomic divergence may indicate that sulfide tolerance is highly polygenic, with shared allele frequency shifts present at many loci with small effects along the genome. Alternatively, H_2_S tolerance may involve substantial genetic redundancy, with non-convergent lineage-specific variation at multiple loci along the genome underpinning similar changes in phenotypes and gene expression. Overall, we demonstrate variability in the extent of convergence across organizational levels and highlight the challenges of linking patterns of convergence across scales.

## Introduction

Evolutionarily independent lineages often evolve similar solutions when facing similar ecological conditions. Although such patterns of convergence may provide evidence for the repeatability and predictability of evolution by natural selection,^1–3^ it is often challenging to determine the extent of convergence across biological levels of organization, and ultimately, whether convergent genomic shifts underpin convergence at other scales.^4^ Convergent evolution is frequently studied in closely related, but geographically disjunct, population pairs exposed to similar sources of divergent selection, providing spatially replicated instances of local adaptation within a clade.^1^ Investigations into convergent evolution in such systems have provided valuable insights across a wide variety of animal and plant systems by highlighting instances of evolutionary convergence at the level of single genes^5,6^ and more polygenic signals of convergence.^7,8^ However, studies that focus on closely related lineages likely biases our understanding of general patterns of convergence, because shared evolutionary responses may be the result of selection on standing genetic variation and constraints associated with shared genomic architecture.^5,9,10^ Similarly, the focus on geographically disjunct populations exposed to seemingly similar selective regimes has the potential to ignore the effects of unquantified environmental variation, which often leads to idiosyncratic evolutionary responses.^11,12^ Studying distantly related taxa that are experiencing a shared selective environment in sympatry eliminates these confounding influences, providing the opportunity to investigate convergence in adaptive strategies and determine the predictability and repeatability of evolution.^13–15^

We took advantage of a unique system to explore shared patterns of adaptive evolution among three sympatric lineages of livebearing fishes (Poeciliidae). In Mexico, populations of *Poecilia mexicana*, *Pseudoxiphophorus bimaculatus*, and *Xiphophorus hellerii* coexist both within the La Gloria sulfide spring complex, which is characterized by high concentrations of naturally occurring hydrogen sulfide (H_2_S), and in adjacent nonsulfidic streams (Figure 1A, B). The presence and absence of H_2_S creates strong divergent selection across habitat types because of its toxic properties. H_2_S binds to cytochrome c oxidase (COX) of the electron transport chain and interrupts oxidative phosphorylation (OxPhos) in mitochondria, ultimately stopping aerobic ATP production necessary for the maintenance of cellular function.^16,17^ At the molecular level, the clear biochemical and physiological consequences of H_2_S predict adaptive modification of genes associated with OxPhos and enzymatic H_2_S detoxification, which is mediated by the mitochondrial sulfide:quinone oxidoreductase (SQR) pathway.^16^ Besides the presence and absence of H_2_S, the two aquatic habitats differ in other physical and chemical attributes, with the sulfide spring having lower dissolved oxygen concentrations and pH, but higher specific conductivity (Figure 1C).

**Figure 1.**
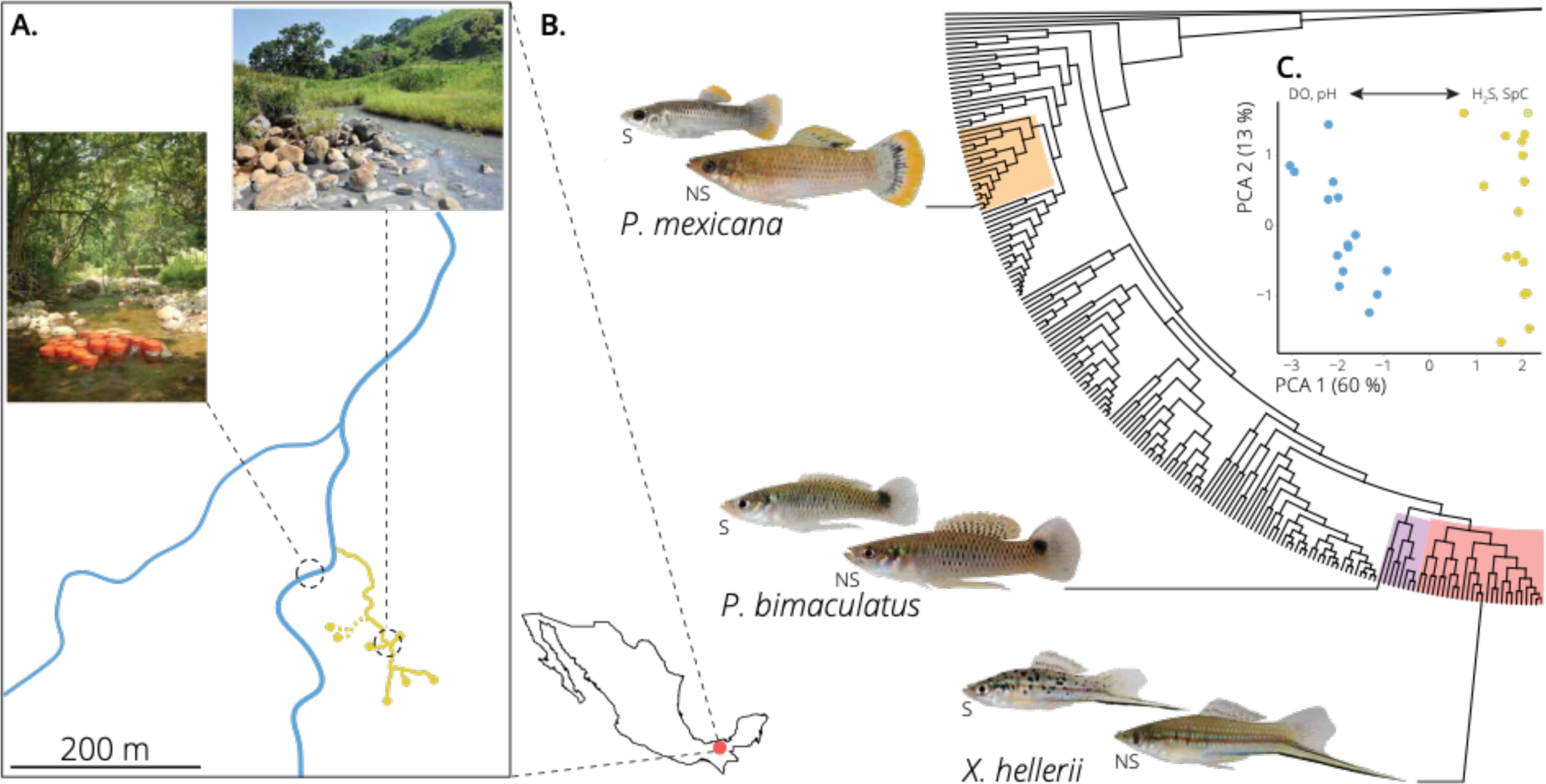
Sympatric populations of three highly divergent poeciliid lineages co-occur in a sulfide spring and adjacent nonsulfidic stream in southern Mexico. A. Map of the study site, where yellow points indicate the H_2_S spring heads of the La Gloria spring system, while yellow lines indicate H_2_S-rich stream segments. Blue lines indicate the adjacent nonsulfidic stream system, Arroyo Caracol. Inset photos depict representative habitats within the two streams (top: La Gloria, sulfidic; left: Arroyo Caracol, nonsulfidic). B. Phylogeny of the family Poeciliidae highlighting the three focal genera: *Poecilia* (orange), *Pseudoxiphophorus* (purple), and *Xiphophorus* (red). Each species’ position within its genus is denoted by the extended branch leading to representative photographs of a sulfidic (S; top) and nonsulfidic (NS; bottom) male of each species. C. Nonsulfidic and sulfidic habitats differed significantly in physical and chemical water parameters. Principal component analysis indicated that sites within the La Gloria spring complex (yellow) and Arroyo Caracol (blue) separated along the primary axis of variation, with nonsulfidic sites exhibiting higher dissolved oxygen concentrations (DO) and pH and sulfidic sites exhibiting higher H_2_S concentrations and specific conductivities (SpC).

The La Gloria system is particularly interesting because it allows for the juxtaposition of evolutionary responses to H_2_S in distantly related species that occur in sympatry with evolutionary responses previously documented in geographically disparate populations of sulfide spring fishes within the family Poeciliidae.^18^ Throughout the Americas, highly endemic poeciliids locally-adapted to sulfide springs have evolved from ancestral populations in adjacent freshwater habitats.^18–20^ Evolutionary responses to the toxic conditions have primarily been studied in geographically replicated populations of *Poecilia mexicana*, which exhibit convergence in morphology, physiology, reproductive life histories, gene expression, and the establishment of reproductive isolation as a byproduct of adaptation via ecological speciation.^18^ While some convergence appears to result from *de novo* modification of the same genes or biochemical pathways (*e.g.*, in OxPhos), there is also evidence for selection on standing genetic variation (*e.g.*, in the SQR pathway).^21–26^ As in other systems, the focus on convergence across geographically replicated lineages within *P. mexicana* may have biased the interpretation of the repeatability and predictability of adaptive evolution in sulfide springs. Indeed, a recent broad-scale study of adaptation to sulfide springs across the Poeciliidae found evidence for convergence in gene expression profiles of ten evolutionarily independent sulfide-spring lineages, but only limited evidence for repeatable patterns of molecular evolution outside of a few mitochondrially-encoded OxPhos genes.^25^ The degree to which this breakdown of molecular convergence at a broader phylogenetic scale is related to differences in the selective regimes among geographically disparate sulfide springs or attributable to lineage-specific evolutionary solutions to a shared source of selection remains unresolved, though evidence suggests that environmental similarity drives the degree of convergence at the transcriptomic level.^27^

In this study, we aimed to identify signatures of convergence associated with the adaptation of three independent lineages to the same H_2_S-rich spring. This approach helped reduce the confounding effects of shared ancestral standing genetic variation and allowed us to stringently test for shared adaptive responses across biological levels of organization in different species. Our results also provided the opportunity to identify possible links between different aspects of convergence across scales, including whether convergence in morphology and gene expression are explicitly underpinned by convergent genomic variation. To address this overarching hypothesis, we (i) identified signatures of convergent evolution in phenotypic traits that have previously been associated with adaptation to sulfide spring environments (body shape, H_2_S tolerance, and gene expression), (ii) established evidence for local adaptation between sulfidic and nonsulfidic populations of each species, and (iii) identified convergent patterns of genomic differentiation among population pairs in sulfidic and nonsulfidic habitats. Overall, we found evidence for convergence in adaptation across multiple levels of organization, but convergence at the genomic level was limited and, outside of the mitochondrial genome, not associated with genomic regions predicted to respond to selection from H_2_S.

## Results

### Phenotypic variation among habitats

Mechanisms of adaptation to H_2_S-rich environments have been studied extensively in poeciliid fishes, including the biochemical and physiological modifications that directly mitigate H_2_S toxicity^25^ and modifications of other traits that are shaped by sources of selection that are correlated with the presence of H_2_S (*e.g.,* hypoxia, increased salinity, changes in trophic resource use, competition, and predation.^18,28^ We quantified variation of natural populations in three complex traits that typically show adaptive divergence upon sulfide spring colonization: body shape, H_2_S tolerance, and gene expression. We found that sulfide spring populations of all three focal species exhibit convergent patterns of trait divergence from adjacent nonsulfidic populations in response to colonization of sulfide springs.

Geometric morphometric analyses of wild-caught, adult fish revealed that body shape significantly differed between sulfidic and nonsulfidic populations of all three lineages (Procrustes ANOVA; species × habitat: *P* < 0.001, *Z* = 7.565, Table S1). Furthermore, there was a significant signal of morphological convergence across the sulfide spring populations of the three species (habitat: *P* = 0.018, *Z* = 2.093, Table S1). Investigation of thin-plate spline transformation grids visualizing shape differences for both the “species × habitat” and “habitat” terms indicated that the shared aspects of variation in body shape between sulfidic and nonsulfidic populations of each species involved divergence in head size, with fish from the sulfide spring exhibiting larger heads than those from the nonsulfidic habitat (Figure 2A).

**Figure 2.**
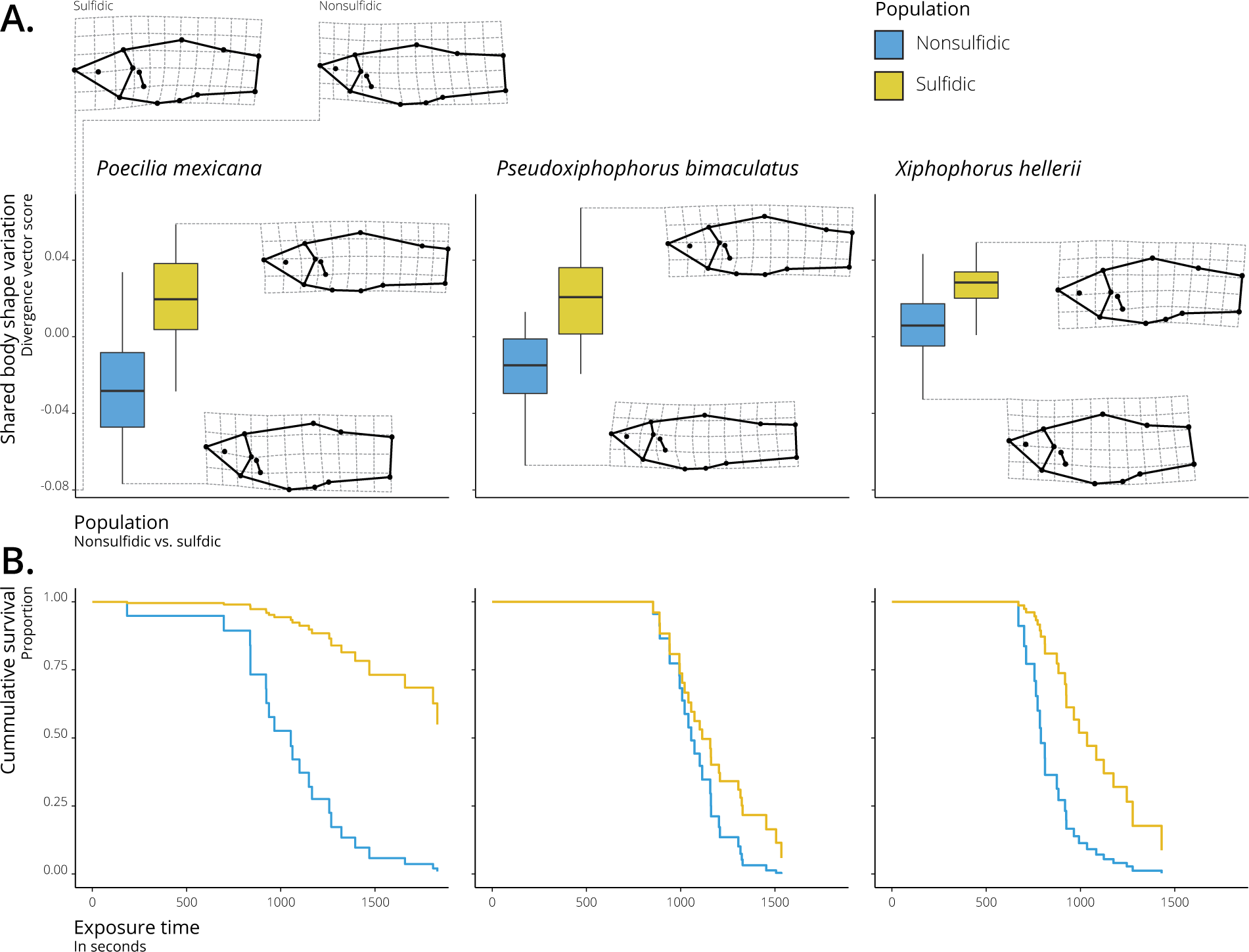
Phenotypic convergence in morphological and physiological traits of sympatric sulfidic populations. A. Convergent morphological evolution among species between habitat types. Body shape differentiation along a shared axis of divergence (habitat type) is depicted by the transformation grids on the top. Significant differences in body shape between populations of each species is depicted in insets (Table S1). B. Convergence in H_2_S tolerance among all populations from the sulfidic habitat. Cumulative survival in acute H_2_S toxicity trials revealed significant differences between populations, which were most pronounced in *P. mexicana* (Figure S1).

Survival analyses indicated that wild-caught, adult fish from all three sulfide spring populations exhibited an increased ability to tolerate acute H_2_S exposure relative to nonsulfidic populations (Figure 2B; Cox regression; habitat: *P* = 0.034, Figure S1). While H_2_S tolerance was clearly affected by habitat of origin, differences in tolerance between sulfidic and nonsulfidic populations also varied among the three species, with sulfidic *P. mexicana* having a higher H_2_S tolerance than both nonsulfidic *Poecilia* and sulfidic populations of the other species (Figure S1). This observation matched microhabitat uses by the three species within the sulfide spring complex; *P. mexicana* was the only species inhabiting immediate spring outlets with the highest H_2_S concentrations, and the other two species were more common in areas with more oxygen-rich water (Figure S2).

Analysis of genome-wide gene expression in gill tissues from wild-caught sulfidic and nonsulfidic fish revealed an overall phylogenetic signal in gene expression variation, with individuals clustering by species and, secondarily, by habitat type (Figure 3A). There were many differentially expressed genes (negative binomial regression, FDR < 0.05) between the sulfidic and nonsulfidic populations of each species, representing 13.4% of all measured genes for *Pseudoxiphophorus*, 20.4% for *Poecilia*, and 20.9% for *Xiphophorus* (Figure 3B). While most gene expression differences were species-specific (Table S2), 245 genes (1.3% of all analyzed genes; 6.3–9.8% of differentially expressed genes in each species) exhibited convergent expression changes across all species (153 upregulated, 92 downregulated; Figure 3B, Table S2), significantly more than predicted by chance (*SuperExactTest*^29^: *P* < 0.0001). As predicted by the biochemical and physiological effects of H_2_S, analysis of Gene Ontology (GO) IDs associated with differentially expressed genes revealed that shared upregulated genes were significantly enriched (*FDR* < 0.05) for associations with mitochondria and biological processes involved in H_2_S toxicity and detoxification, including aerobic respiration and the electron transport chain (OxPhos), enzymatic H_2_S detoxification, and the processing and transport of sulfur compounds (Table S3). Notably, we found consistent upregulation of genes associated with the primary target of H_2_S toxicity (COX) and the primary H_2_S detoxification pathway (SQR pathway; Figure 3C–E). There was no evidence for significant enrichment of any GO terms in downregulated genes (Table S3).

**Figure 3.**
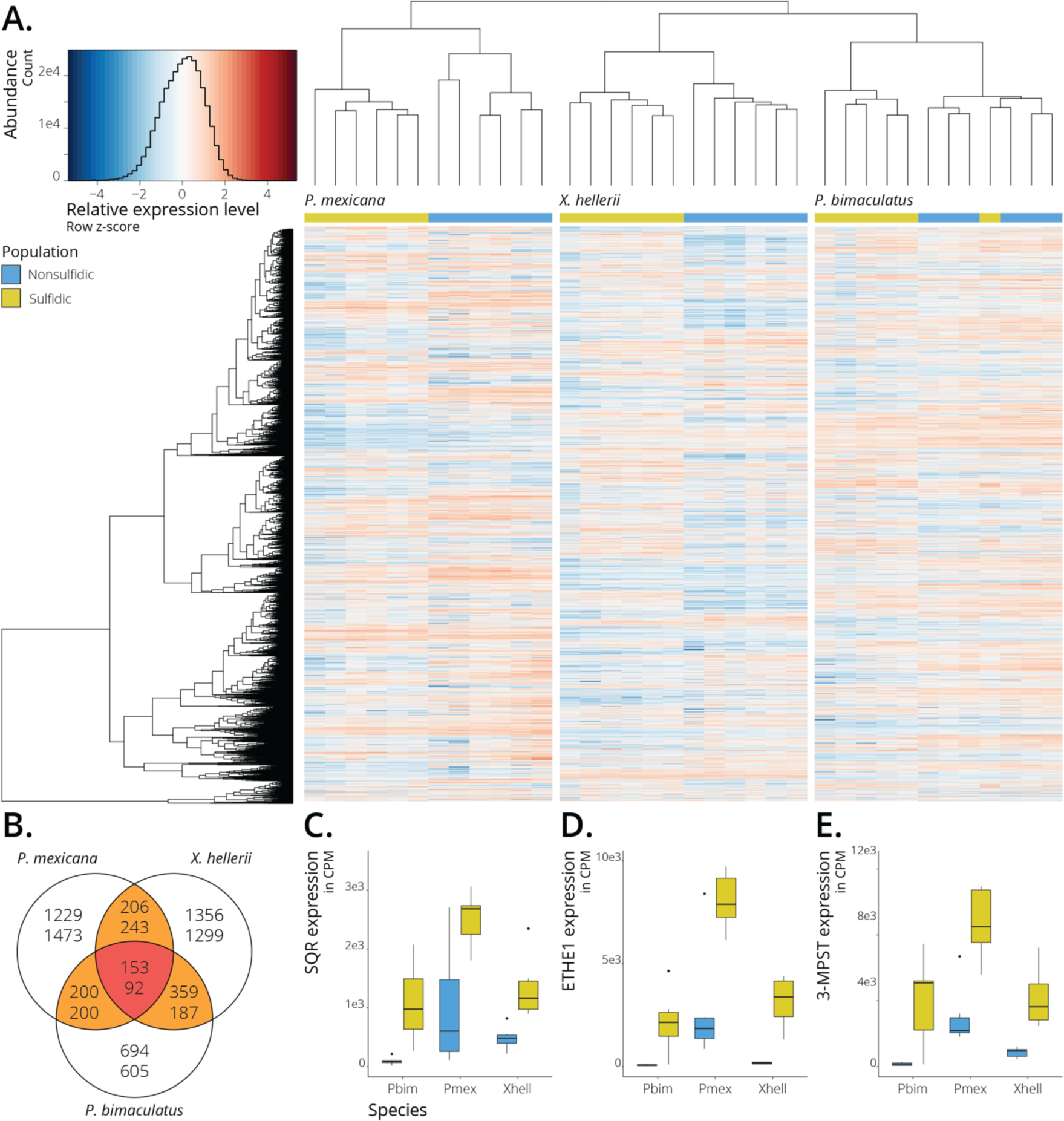
Convergent shifts in gene expression underlie local adaptation. A. Hierarchical clustering analysis indicates that gene expression profiles cluster by habitat type within each species. Each row represents expression data for one gene, and each column corresponds to an individual. Genes with higher relative expression in individuals from sulfidic habitats are indicated in red, genes with lower relative expression in blue. B. Venn diagram showing the number of differentially expressed genes between the sulfidic and nonsulfidic populations of each species and those shared among species. Top numbers are genes upregulated in the sulfidic habitat; bottom numbers are downregulated genes. C-E. Sulfidic populations consistently exhibit higher expression of key H_2_S detoxification genes (SQR: sulfide:quinone oxidoreductase; ETHE1: persulfide dioxygenase; 3-MPST: 3-mercaptopyruvate sulfurtransferase) than nonsulfidic populations of the same species. Box plots of counts per million (CPM) by population. All boxplots have the following elements: center line, median; box limits, upper and lower quartiles; whiskers, ×1.5 interquartile range; points, outliers closed circles.

### Population Structure

As phenotypic divergence between sulfidic and nonsulfidic populations of each species was measured using wild-caught fish, we cannot rule out the possibility that both adaptation and phenotypic plasticity contribute in some part to phenotypic responses to the colonization of the sulfidic habitat. However, since divergent selection on key ecological traits generally reduces gene flow between populations, local adaptation and phenotypic plasticity are predicted to have different population genomic signatures. When plasticity alone underpins phenotypic differentiation across environmental contrasts, we expect to see little genome-wide population structure, as gene flow between habitats prevents genomic differentiation from establishing. In contrast, where divergent selection has resulted in the establishment of barriers to gene flow, we expect to find lower rates of gene flow between populations and signals of population structure, even though the La Gloria sulfide spring complex is spatially restricted (∼200 m in length), flows directly into the adjacent nonsulfidic stream (Figure 1A), and there are no physical barriers preventing fish movement between habitat types. Additionally, previous investigations into the signatures of gene expression associated with adaptation of *P. mexicana* to sulfide springs have provided evidence of heritable variation in gene expression,^30,31^ which highlights the potential for a component of adaption to be underpinned by genomic variation rather than plasticity alone. To test for genetic signatures of local adaptation, we sampled 20 individuals per species in each habitat type and conducted population genomic analyses based on whole-genome resequencing (average sequence depth per individual: 3.8 ± 0.4; Table S4) to infer population structure between the sulfidic and nonsulfidic populations. Maximum-likelihood-based estimates of admixture proportions with two inferred clusters (*k* = 2) unambiguously clustered individuals by their habitat type of origin and only detected low levels of admixture between sulfidic and nonsulfidic populations within each species (Figure 4A), indicating a substantial reduction in gene flow between the two habitats. Principal component analysis (PCA) corroborated these results, consistently separating populations by habitat type along the primary axis of genetic variance in each species (17.5–53.5% of genomic variation explained by PC1; Figure S3). Both structure analysis and PCA identified two *P. mexicana* individuals that appear to be early generation hybrids (F1 and a possible F2 backcross), while all other individuals of all species exhibit clear assignment to their habitat of origin with limited, asymmetric gene flow from sulfidic into nonsulfidic populations.

**Figure 4.**
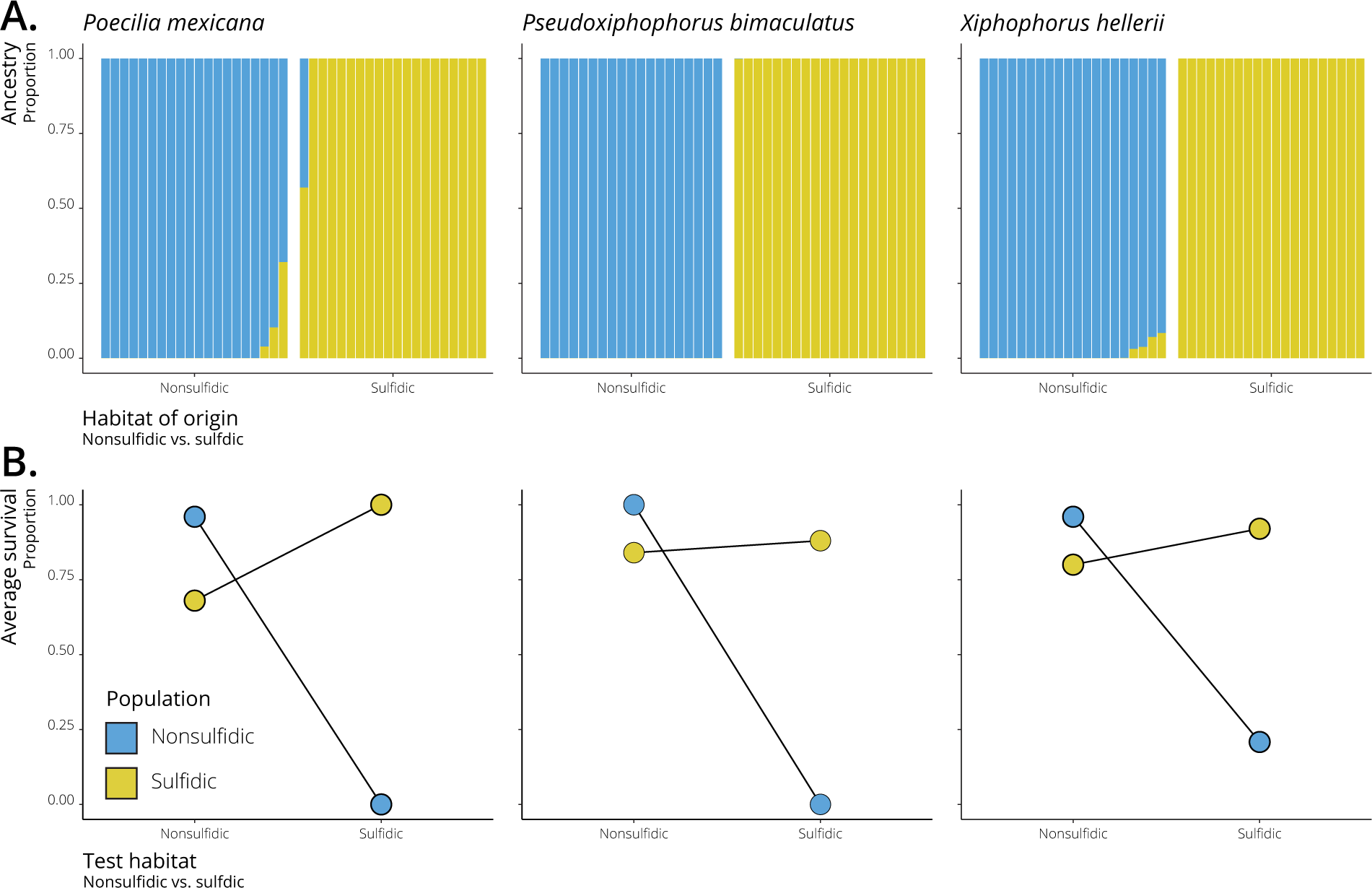
Nonsulfidic and sulfidic populations of three sympatric species exhibit limited gene flow between habitats mediated by strong natural selection against migrants. A. Admixture analyses revealed substantial population genetic differentiation in each population pair. Individuals are grouped by sampling site, corresponding with the detected genetic population clusters (*K*=2) shown in blue (nonsulfidic) and yellow (sulfidic). B. Translocation experiments revealed strong selection against migrants, especially those from nonsulfidic to sulfidic habitats (Table S5). Fish were transferred between sites (test habitat) to estimate survival of migrants between habitats, with individuals collected from the nonsulfidic population in blue and from the sulfidic population in yellow.

We also conducted *in situ* reciprocal translocation experiments of wild-caught, adult fish, which explicitly tested for local adaptation^32^ and natural selection against migrants.^33^ We found strong, habitat-specific differences in the survival of sulfidic and nonsulfidic populations of each lineage (Figure 4B; binomial GLMM; habitat of origin × testing habitat: *P* < 0.001; Table S5). Fish tested in their own habitat had significantly higher survival rates (88–100%) than those tested in the opposite habitat type, with the lowest survival observed for fish moved from the nonsulfidic into the sulfidic habitat (0–21%). Fish moved from the sulfidic into the nonsulfidic habitat had comparatively high survival (68–84%)—though significantly lower than resident nonsulfidic fish—ultimately supporting the observed asymmetry of gene flow between habitat types. Overall, this experiment supported the inference of local adaptation and that natural selection against migrants likely serves as a strong mechanism generating population structure between sulfidic and nonsulfidic populations.

### Landscape of Genomic Divergence

The explicit biochemical and physiological effects of H_2_S—along with the convergent changes at the transcriptome level—predict clear targets of selection at the genomic level. We first explored potential convergent changes in the mitochondrial genome, which includes genes for three subunits of H_2_S’ primary toxicity target (COX). Mitochondrial haplotype networks revealed that sulfidic populations harbor mitochondrial lineages that are highly divergent from those in the nonsulfidic populations (Figure 5A). Consistent with the asymmetric patterns of gene flow, we found sulfidic mitochondrial haplotypes in nonsulfidic habitats, especially in *P. mexicana* and *X. hellerii*, while nonsulfidic mitochondrial haplotypes were not found in sulfidic populations (except for one apparent first-generation hybrid in *P. mexicana* described above). Using analyses of molecular evolution, we detected evidence for positive selection (elevated nonsynonymous to synonymous substitution rates, μ; likelihood-ratio test, *P* < 0.01; Table S6) acting on five mitochondrial genes in all three sulfidic populations, including two COX subunits (COX1 and COX3) and genes encoding subunits of OxPhos complexes I (ND2 and ND3) and III (CYTB), corroborating previous analyses of sulfide spring fishes.^21,25^

**Figure 5.**
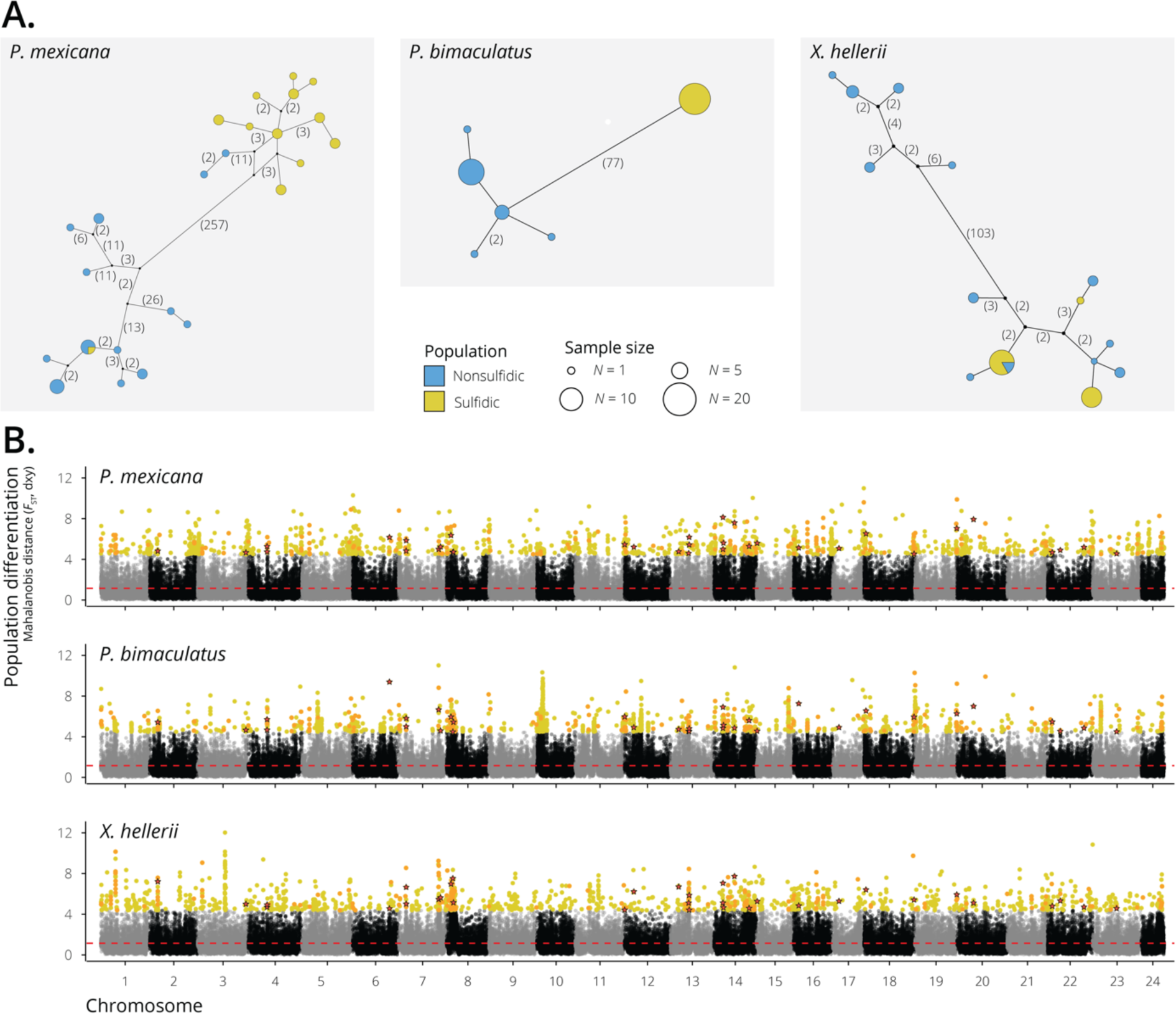
Convergent patterns of genomic adaptation are limited to the mitochondrial genome and few nuclear loci. A. Haplotype networks based on concatenated mitochondrial genes indicate high levels of mitochondrial divergence between nonsulfidic (blue) and sulfidic (yellow) populations, with asymmetrical introgression of sulfidic mitochondria into nonsulfidic individuals. The relative size of each circle indicates the number of individuals with a particular mitochondrial haplotype, while numbers in parentheses represent the total number of nucleotide differences between haplotypes. B. Mahalanobis distance (–log_10_ *P*-values of *F*_ST_ and *d*_XY_) of non-overlapping 5-kb windows across the genome as a measurement of genomic divergence driven by natural selection. Outlier windows (top 1%) unique to each population pair are shown in yellow, while those shared between two population pairs are in orange. The outlier windows shared among all three population pairs are denoted with red stars.

Unlike in the mitochondrial genome, recombination in the nuclear genome can mediate highly heterogeneous divergence during local adaptation. When few genomic regions are directly involved in adaptation, these regions often exhibit disproportionally high divergence (genomic islands of divergence) against a background of regions of low divergence that is homogenized by gene flow.^34,35^ If the observed convergent morphological and transcriptomic differentiation had a convergent genomic basis across lineages, we would predict overlapping genomic islands of divergence between sulfidic and nonsulfidic populations across the three species, including regions with mitochondrially-functioning genes that are involved in H_2_S toxicity and detoxification. Considering the evidence for positive selection on mitochondrially encoded genes, we also expected to detect evidence for mito-nuclear coevolution,^36^ with disproportionally high divergence in the nuclear genes encoding subunits of OxPhos complexes I, III, and especially IV. To determine if selection on the same genomic regions underlies convergence to sulfide spring environments, we used the whole-genome resequencing data described above to characterize the landscape of genomic divergence in each population pair using a combination of relative (*F*_ST_) and absolute (*d*_XY_) divergence metrics in non-overlapping 5-kb windows.^37,38^ Global estimates of relative divergence varied considerably among species (mean *F*_ST_ = 0.23 for *Poecilia*, 0.06 for *Pseudoxiphophorus*, and 0.03 for *Xiphophorus*; Figure S4), while absolute divergence across the genome was similar (mean *d*_XY_ = 0.003 for *Poecilia*, 0.004 for *Pseudoxiphophorus*, and 0.003 for *Xiphophorus*; Figure S5).

Using a composite analysis that simultaneously considered relative and absolute divergence,^39^ we observed heterogeneous divergence across much of the genome within and among population pairs, rather than locally pronounced and shared genomic islands of divergence (Figure 5B; Table S7). By identifying the most differentiated regions between sulfidic and nonsulfidic individuals within each species (top 1% most differentiated windows; 1,341–1,366 5-kb windows per species), we found that regions of elevated divergence were significantly enriched for genes involved in oxygen transport (*Pseudoxiphophorus*) and immune function (*Xiphophorus* and *Poecilia*), including hemoglobins and major histocompatibility complex genes (GO enrichment analysis: *FDR* < 0.05; although in *Poecilia* this enrichment was non-significant following FDR correction; Tables S8). To test for convergent patterns of differentiation, we identified regions that were amongst the most differentiated windows across all three species (top 1% most differentiated in each of the three comparisons to account for variation in average genome-wide differentiation across the three comparisons; Figure 5B). Thirty-four strongly differentiated regions—containing 28 genes—were shared across all three species (significantly more than expected by chance; *SuperExactTest*, *P* < 0.0001), and 102–156 of these regions were shared between pairs of the three species (Table S7). There was no significant enrichment of GO terms in strongly differentiated regions shared among all lineages, and we found no evidence for genes with known roles in H_2_S detoxification or processing in regions of elevated divergence within species (Tables S8). We did observe some overlap of shared differentiated regions and differentially expressed genes within each species (150–198 genes), but not more than would be expected by chance (Fisher’s Exact Test, *P* = 0.24–1.0).

We also carried out site-frequency-spectrum based scans for selective sweeps^40^ in each sulfidic population and compared putative sweep regions across the three sulfidic and nonsulfidic contrasts. We found no shared selective sweep outlier regions across all three species (Figure S6), and the small subset of genes shared among pairs of species was not enriched for any biological functions (Table S9). *Pseudoxiphophorus bimaculatus* exhibited enrichment of both ion transport and oxygen transport genes, as well as evidence for a selective sweep in the region containing SQR, a key H_2_S detoxification enzyme (Table S10). However, neither *P. mexicana* nor *X. hellerii* exhibited any evidence for selective sweeps in regions containing genes related to H_2_S, and only *X. hellerii* exhibited limited enrichment for GO terms in sweep regions (plasma membrane associated genes; Table S10).

## Discussion

The three poeciliid fishes that have colonized the La Gloria sulfide spring complex provided a unique opportunity to examine how distantly related lineages respond to the same sources of divergent selection. This approach helped reveal which aspects of evolutionary responses are repeatable and predictable and how responses and the predictability of these responses vary across biological scales of organization. We found evidence for local adaptation in each of the lineages and detected signatures of convergence in phenotypic traits that span levels of organismal organization. In addition, there was convergent selection on mitochondrially-encoded genes associated with H_2_S toxicity. However, evidence for convergence in the nuclear genomes was scant and not associated with genomic regions predicted to respond to selection from H_2_S. Hence, landscapes of genomic divergence can be highly idiosyncratic even when species experience the same selective regime in sympatry, suggesting that evolutionary outcomes are contingent on the genomic substrates that selection is acting on. These results also highlight the potential methodological challenges of identifying convergent shifts associated with highly polygenic traits, phenotypic traits that are associated with high genetic redundancy, or traits underpinned by convergent structural variation.

### Evidence for Convergence in Traits Associated with Adaptation to Sulfide Springs

Consistent with previous findings,^41,42^ we found significant differences in body shape between sulfidic and nonsulfidic populations of all three lineages at La Gloria, with sulfide spring fishes exhibiting larger heads than those in nonsulfidic habitats. Previous studies have indicated that increased head size in sulfide spring poeciliids reflects a response to selection on oxygen acquisition in the hypoxic sulfide springs, as head size is correlated with increased gill surface area and higher ventilation capacity in sulfide spring populations of *P. mexicana*^41,43^ and other fishes.^44,45^

Convergent phenotypic differentiation in H_2_S tolerance between sulfidic and nonsulfidic fishes was also observed. Sulfide spring populations exhibited higher tolerance to acute H_2_S exposure than populations from adjacent nonsulfidic populations, and, accordingly, nonsulfidic fishes suffered high mortality when placed into the natural conditions of the sulfide spring. Previous studies have demonstrated that sulfide spring populations of *P. mexicana* consistently exhibit higher tolerance to acute exposures to H_2_S, whether it is quantified at a molecular or whole-organism level.^25,41^ Increased tolerance results from modification of the expression and protein coding sequences of H_2_S toxicity and detoxification genes in well characterized biological pathways.^21,25^ Additionally, our findings supported those of previous studies carried out in more divergent poeciliid lineages occupying sulfide springs across the Neotropics, which indicated that convergent shifts in the expression of H_2_S toxicity and detoxification genes is correlated with H_2_S tolerance.^25,46^ We found that upregulated genes in the gills of sulfidic populations from all three species were enriched for genes with mitochondrial functions and H_2_S processing, including genes (*COX* and *SQR*) demonstrated to exhibit adaptations for processing H_2_S or resisting its toxicity in sulfide spring *P. mexicana*.^21,25^ Ultimately, these shared expression patterns across the three sulfide spring populations provide robust support for convergent regulation of genes with known roles in H_2_S toxicity and detoxification.

While these signals of convergence are substantial, we cannot eliminate the possibility that differences in body shape, sulfide tolerance, and gene expression observed in our experiments with wild-caught specimens were at least in part shaped by plastic responses to H_2_S exposure, rather than evolved differences between populations. Many physiological traits, such as sulfide tolerance and gene expression, are known to exhibit short-term plastic responses to changes in environmental conditions.^30^ Common-garden rearing of these species will be needed to better understand how genetic differentiation and plasticity interact to shape gene expression patterns observed in nature; however, laboratory rearing of these sulfidic populations has proven difficult (Greenway and Tobler, personal observation). Previous common-garden experiments of other sulfidic and nonsulfidic *P. mexicana* populations, notably less divergent than the populations examined in this study, have provided evidence for a heritable basis to divergence in body shape,^47^ sulfide tolerance,^48^ and expression variation in H_2_S-processing and OxPhos-related genes.^30,31^ Furthermore, our population genetic analyses demonstrated that fish collected from the sulfidic and nonsulfidic habitat exhibit very little gene flow between them, despite the possibility of panmixia due to the connectedness of the streams in which they are found, suggesting that the phenotypic differentiation we observed likely does not solely reflect habitat-specific inducible effects but are at least in part the result of evolved differences.

### Genetic Basis of Adaptation

Even with clear evidence for convergent phenotypic responses to shared environmental conditions, we found that patterns of convergence were not consistent across scales of biological organization. Specifically, we found limited evidence for convergent evolution at the genomic level, suggesting that convergence in morphology, physiological tolerance, and gene expression do not simply stem from genomic differentiation at one, or few, genomic loci. Our results show that nuclear genes typically implicated in H_2_S adaptation were not associated with shared or species-specific outlier regions, including genes of the SQR pathway that are involved in enzymatic H_2_S detoxification and nuclear genes associated with COX and other OxPhos components. These results stand in stark contrast to previous investigations of replicated *P. mexicana* populations adapted to sulfide springs across southern Mexico, where a variety of approaches have demonstrated predictable evolution of genes with known roles in H_2_S detoxification and metabolism.^21,23,24^ This high level of convergence among closely related populations paired with the limited convergence observed among divergent lineages in the present study suggests that even the strong physiological demands of H_2_S toxicity are not sufficient to constrain adaptation to one or a few solutions at the genomic level. Rather, access to shared pools of genetic variation, as is the case for replicated *P. mexicana* populations,^23,26^ appears to facilitate convergence at the genomic level. These results align with previous studies investigating genome-wide patterns of changes in protein coding sequences (based on transcriptomic data), which found little evidence for convergent evolution of protein coding sequences outside of the mitochondrial genome.^25^ However, in line with previous studies, we did observe consistent modification of genes encoded by the mitochondrial genome across all three sulfidic populations, including the direct targets of H_2_S toxicity (*COX* subunits). Additionally, nonsulfidic mitochondrial haplotypes were not found in individuals sampled from the sulfidic habitat (except for one apparent F1 hybrid), suggesting a critical role of mitochondrial DNA in mediating survival in the toxic conditions. Despite evidence for selection on mitochondrial genes,^49^ it remains unclear how genetic variation in mitochondrial versus nuclear loci contribute to adaptive variation in OxPhos function. The evolution of H_2_S-resistance could be largely driven by modification of mitochondrial *COX* subunits that form the reactive center of the protein and without coevolutionary changes in the corresponding nuclear subunits. This finding challenges the general paradigm that adaptation of mitochondrial function coincides with mito-nuclear coevolution.^49^

While evidence of selection on nuclear genes associated with H_2_S detoxification and OxPhos was sparse, functional annotation of shared and unique outlier regions did find limited evidence for enrichment of genes involved in oxygen transport and immune function (Tables S8). Evidence for selection on oxygen transport proteins, such as hemoglobin is not surprising, as H_2_S is capable of binding to and impairing hemeprotein function.^50,51^ The presence of hemoglobin genes in outlier regions also confirms findings of positive selection on hemoglobin genes in other sulfide spring poeciliids.^52^ In contrast, the large number of genes associated with immune function was not expected, although immune genes are known hotspots for selection in vertebrates.^53^ Selection on immune genes could be driven by differences in parasite and pathogen communities between sulfidic and nonsulfidic habitats, which have been documented in another sulfide spring inhabited by a poeciliid fish.^54^ Alternatively, there have been long-standing, but poorly investigated, hypotheses about the potential role of H_2_S-oxidizing microsymbionts in adaptation to sulfide spring environments,^16,55^ and changes in host-microbe interactions upon sulfide-spring colonization may also exert selection on immune genes.^56^

Even though H_2_S has clear-cut biochemical and physiological effects, tolerance to this toxicant involves numerous genes. OxPhos complexes alone contain over 80 nuclear-encoded genes,^57^ and dozens of genes contribute to the physiological processing of H_2_S and other sulfur compounds.^58,59^ For this reason, the lack of convergence we observed at the genome level could be the result of the polygenic nature of H_2_S tolerance, which is known to pose substantial challenges for detecting patterns of genomic convergence, since associated loci are unlikely to form large islands of divergence but rather be scattered throughout the genome. It is possible that subtle allele frequency shifts across multiple polymorphic OxPhos or H_2_S processing genes could result in adaptive modifications to these pathways without leaving a molecular signature detectable by scans for genomic differentiation between divergent species pairs.^60,61^ Such subtle allele frequency shifts are now detectable when closely related populations with shared allelic variants are exposed to similar sources of selection^62^ but not in highly divergent population pairs that were investigated here. It is also important to emphasize that environmental gradients between sulfidic and adjacent nonsulfidic habitats are complex (Figure 1C). The multifarious sources of selection have caused adaptive modifications of whole suites of complex traits beyond H_2_S tolerance, and most of these traits likely have different genomic bases.^18^ Finally, another important factor influencing our ability to detect genomic convergence is whether H_2_S tolerance involves substantial genetic redundancy, where many different combinations of alleles throughout the genome can underpin the same phenotype. If genetic redundancy is high, lineage-specific responses to selection that are in part shaped by historical contingencies^63–65^ may underpin the convergence signatures of phenotypic traits and gene expression without genomic convergence across species.^66^

It is also possible that adaptation to H_2_S may primarily be driven by changes in mitochondrial protein function and the expression of detoxification genes, with lineage-specific mechanisms producing the convergent gene expression patterns uncovered by transcriptome analyses. Because differentially expressed genes were not overrepresented in the outlier genome regions as might be expected by selection acting on *cis*-regulatory elements,^24^ outlier regions may harbor *trans*-regulatory elements mediating adaptive shifts in gene expression.^67^ Divergent selection acting on regulatory elements has been implicated in adaptive divergence for several well-known models in evolutionary ecology^68–70^. The problem of detecting the genetic basis of trait variation by any of the above means can be exacerbated when adaptive evolution gives rise to reproductive isolation. Selection on small-effect loci spread across the genome can facilitate strong and stable reductions in gene flow over short timescales,^71,72^ similar to the reduced gene flow observed for all three lineages in our study. High levels of background divergence resulting from post-speciation divergence can further obscure the signatures of selection on the loci mediating adaptation during earlier stages of divergence.^73–75^

## Conclusions

Our results showed that while convergent evolution in response to shared selective pressure is present across diverse lineages, patterns of convergence to extreme environments are not consistent across levels of biological organization. We also demonstrated that mechanistically linking strong signals of convergence in morphology, physiology, and gene expression to genomic variation is challenging due to the magnitude of ways in which different aspects of genomic variation interplay to affect phenotypic variation. Although we observed signatures of independent local adaptation and reproductive isolation between sulfidic and nonsulfidic populations in three highly divergent lineages, variation in the complexity and genomic architecture of traits and the degree of genetic redundancy may reduce our ability to confidently identify signals of convergence at the genome level. While substantial progress has been made in understanding the molecular basis of trait evolution, methodological and statistical limitations are prevalent when traits are complex, leading to an underrepresentation of studies covering the adaptation of non-model organisms to complex ecological stressors.

## STAR Methods

### Study sites and sample collection

In the foothills of the Sierra Madre de Chiapas, several freshwater springs rich in naturally occurring H_2_S form the La Gloria spring complex.^41,76^ The spring complex is located near the city of Teapa, Mexico, and is part of the Río Pichucalco drainage. Habitats with high H_2_S concentrations are spatially restricted (∼200 m in length), and the spring run flows directly into a nearby nonsulfidic stream (Arroyo Caracol). There are no physical barriers that prevent fish movement between the sulfidic spring run and the nonsulfidic stream, but there are stark differences in water chemistry (Figure 1). Populations of the poeciliid fishes *Poecilia mexicana, Pseudoxiphophorus bimaculatus*, and *Xiphophorus hellerii* have colonized H_2_S-rich habitats at La Gloria, and ancestral populations also occur in adjacent nonsulfidic habitats.^77^ Note that the population of *P. mexicana* at La Gloria likely belongs to an endemic species described from sulfide springs, *P. sulphuraria.*^77,78^ All samples used in this study were collected by seine from the La Gloria sulfide spring complex and nonsulfidic habitats in the adjacent Arroyo Caracol. The sole exception were *Pseudoxiphophorus* gill tissues for the nonsulfidic population, which were collected from another nearby stream (Table S11).

Physical and chemical water parameters were analyzed for several sites within the La Gloria spring complex and Arroyo Caracol. Temperature, specific conductivity, pH, and oxygen content were measured using a Hydrolab Multisonde 4A (Hach Environmental). Environmental H_2_S concentrations were measured with a methylene blue assay using a Hach DR1900 Portable Spectrophotometer (Hach Company, Loveland, CO, USA). Measurements and calibration of probes were conducted according to the manufacturer’s recommendations. At least three Hydrolab readings and two H_2_S samples were taken at each site. Measurements were averaged for each site prior to use in a principal component analysis.

### Evidence for convergence in traits associated with adaptation to sulfide springs

To identify potential convergent evolution across the three focal species, we investigated three distinct complex traits that have previously been implicated in adaptation to sulfide spring environments: (1) body shape; (2) tolerance to acute H_2_S exposure; and (3) genome-wide patterns of gene expression.

#### Analysis of body shape

To test for divergence in body shape among populations in sulfidic and nonsulfidic habitats, adult *Poecilia*, *Pseudoxiphophorus*, and *Xiphophorus* were collected from the two sites using seines. Lateral photographs were taken for all specimens using a Canon EOS 400D digital camera (Canon USA Inc., Lake Success, NY, USA) mounted on a copy stand. We digitized 14 morphological landmarks (Figure S7) on each photograph using the software program tpsDig.^79^ We extracted coordinates of these digitized landmarks to conduct geometric morphometric analyses^80^ using the GEOMORPH package in R.^81^ We first aligned landmark coordinates via least-squares superimposition implemented in the gpagen function to remove effects of translation, rotation, and scale, yielding aligned Procrustes coordinates that describe body shape variation and centroid size as a measure of size. Procrustes coordinates were then analyzed using a Procrustes ANOVA (procD.lm) using residual randomization over 9999 iterations and type III sums of squares. Predictor variables included species (*Poecilia*, *Pseudoxiphophorus*, or *Xiphophorus*), habitat (sulfidic vs. non-sulfidic), and sex as factors, centroid size as a covariate, as well as their interaction terms. *P*-values were based on a Cohen’s f-squared sampling distribution. For data visualization, we calculated divergence vector scores^82^ for each individual based on the first principle component of the among-group covariance matrix for the habitat term from a preparatory MANCOVA using the same model as for the Procrustes ANOVA.^83^ This approach allowed us to visualize aspects of body shape variation associated with habitat type among species (*i.e.*, convergent aspects of body shape), while accounting for all other terms in the model.

#### Tolerance to acute H_2_S exposure

To test for differences in H_2_S tolerance between individuals collected from the sulfidic and nonsulfidic populations, we conducted acute H_2_S exposure trials, subjecting wild-caught fish to logarithmically increasing concentrations of H_2_S.^41^ We collected adult *Poecilia*, *Pseudoxiphophorus*, and *Xiphophorus* from the two sites using seines, transferred them to insulated coolers filled with water from the collection site, and transported them to a nearby field station. Fish were kept in population and species-specific holding tanks with aeration for at least 24 hours prior to testing. To standardize experimental conditions, water from the collection sites was slowly replaced with H_2_S-free well water over the first 8 hours in the holding tanks. Water was continuously aerated and filtered during this time, and the fish received no food.

For the experiment, we prepared stock solutions of 10 mM aqueous H_2_S solution by dissolving 2.4 g sodium sulfide hydrate (Na_2_S·6H_2_O) into 1 L of well water deoxygenated through purging with nitrogen.^84^ Individual fish were placed into clear plastic containers with 150 mL water from the holding tanks and allowed to acclimate for 5 minutes. Following acclimation, 10 mL of H_2_S solution were added to the experimental container at 2-minute intervals using a syringe placed under the water surface. Each fish was observed as H_2_S concentration increased in the experimental container. We measured the time until the fish lost equilibrium, at which point the fish was removed from the container, sexed, weighed, and placed into a heavily aerated recovery tank. Experiments were ended after 32 minutes (15 sulfide additions) if a fish did not lose equilibrium.

We analyzed variation in H_2_S tolerance using a survival analysis (Cox regression) with the coxph function implemented in the R package SURVIVAL.^85^ We used time to loss of equilibrium as the response variable, and individuals that did not lose equilibrium during the experiment were censored. Body mass (log_10_-transformed), species, sex, and habitat of origin (sulfidic vs. nonsulfidic) were used as independent variables. Adjusted survival curves were generated for the habitat term of the Cox regression to visualize expected survival curves, using the ggadjustedcurves function from the SURVMINER package.^86^

#### Transcriptome sequencing and alignment

We used an RNA-sequencing approach to identify differentially expressed genes between the sulfidic and nonsulfidic populations of each species. We collecteded adult female individuals of *Poecilia*, *Pseudoxiphophorus*, and *Xiphophorus* in the La Gloria sulfide spring and from the adjacent nonsulfidic stream. Following capture with a seine, fish were instantly euthanized, and gill tissues were extracted from both sides of the head and immediately preserved in 2 mL of RNAlater (Ambion, Inc.). We selected gill tissue because the gills mediate physiological processes necessary for the maintenance of homeostasis,^87^ are in direct contact with the H_2_S-rich water,^16^ and show strong, heritable transcriptional responses upon H_2_S exposure.^30,46^ RNA was isolated from gill tissues, and cDNA libraries were constructed for each sample as described for samples in previous studies.^25,46^ Briefly, we extracted total RNA from pulverized gill tissues using the NucleoSpin RNA kit (Macherey-Nagel GmbH & Co. KG, Düren, Germany) following the manufacturer’s protocol. We conducted mRNA isolation and cDNA library preparation with the NEBNext Poly(A) mRNA Magnetic Isolation Module (New England Biolabs, Inc., Ipswich, MA, USA) and NEBNext Ultra Directional RNA Library Prep Kit for Illumina (New England Biolabs, Inc., Ipswich, MA, USA), following the manufacturers’ protocol with minor modifications. Each cDNA library was assigned a unique barcode, quantified, and pooled in sets of 11–12 samples that were sequenced together in a way that different species and habitat types were sequenced together. cDNA libraries were sequenced on an Illumina HiSeq 2500 with paired-end, 100-bp reads at the Washington State University Spokane Genomics Core.

Raw RNA-sequencing reads were sorted by barcode and trimmed twice (*--quality* 0 to remove adapters, followed by *--quality* 24) with Trimgalore!^88^ Trimmed reads for all 36 individuals were mapped to the *X. maculatus* reference genome used for genomic analyses^89^ using BWA-MEM.^90^ On average, ∼97% of reads mapped across all individuals (Table S12). We functionally annotated genes from the *X. maculatus* reference genome by extracting the longest transcript per gene (using the perl script gff2fasta.pl^91^) and then searching against the human SWISSPROT database (critical E-value 0.001; access date 11/15/2018) using BLASTx.^92^ Each *X. maculatus* gene was annotated with the top BLAST hit based on the top high-scoring segment pair. Annotations were used for analysis of differentially expressed genes as well as for analyses of outlier regions from genomic data (see below).

We used STRINGTIE v.1.3.3b^93,94^ to quantify the number of RNA-seq reads mapped to each gene in the *X. maculatus* reference genome for each individual and used a Python script provided with STRINGTIE (prepDE.py) to generate a read counts matrix.^94^ We then removed genes that did not have at least two counts per million in 3 or more individuals across all species, resulting in a set of 18,598 genes for analysis of gene expression patterns. We first conducted hierarchical cluster analysis on the full set of retained genes. The hierarchical cluster analysis was performed on log_2_-transformed counts-per-million and visualized with the heatmap.2 function in the GPLOTS package in R.^95^

To identify differentially expressed genes, we used generalized linear models (GLMs) as implemented in the Bioconductor package EDGER.^96^ We used glmFit to fit a negative binomial GLM to the normalized read counts of each gene based on tagwise dispersion estimates and a design matrix describing the comparison between the sulfidic and nonsulfidic population of each species. We assessed statistical significance with the GLM likelihood-ratio test with a false-discovery rate (FDR) of < 0.05 calculated with the Benjamini-Hochberg correction.^97^ We then intersected the significantly upregulated and downregulated genes from all three species-specific comparisons separately to identify genes with consistent evidence for differential expression among all three species. After identifying the set of genes with differential expression between sulfidic and nonsulfidic populations of each species, and genes with evidence for shared differential expression across all three species-specific analyses (Table S2), we used a Gene Ontology (GO) enrichment analysis to explore the putative biological functions of these candidate sets of genes (Tables S3). We first annotated all genes that had a match in the human SWISSPROT database with GO IDs.^98^ In total, 10,817 unique SWISSPROT annotations were associated with a term in the GO database. We then tested for the enrichment of specific GO IDs in these sets of genes with differential expression relative to the full dataset of 18,598 analyzed genes using GORILLA (*P*-value threshold: 0.0001, accessed 04/11/2019).^99^

### Evidence for local adaptation

To test for local adaptation in each of the three sulfidic/nonsulfidic population pairs, we used whole-genome resequencing to assess the contemporary levels of population genetic structure. We also conducted a reciprocal translocation experiment to compare the survival of individuals across the sulfidic/nonsulfidic environmental gradient.

#### Genomic DNA library preparation and whole genome sequencing

We resequenced the genomes of 120 individuals (40 per species, 20 per population). DNA was extracted from fin clip tissues that were stored in ethanol using the NucleoSpin Tissue kit (Macherey-Nagel GmbH & Co. KG, Düren, Germany) following the protocol in the user manual (version January 2017/Rev. 17). For the final elution step, extracted DNA was eluted in 100 µL of elution buffer.

Library preparation and sequencing were conducted at the Washington State University Spokane Genomics Core. Genomic DNA (gDNA) was quantified using the Qubit 2.0 fluorometer with the dsDNA HS assay kit (Thermo Fisher Scientific, Waltham, MA). One hundred nanograms of gDNA were used as input for library preparation using the TruSeq Nano DNA Library Prep Kit (Illumina, San Diego, CA). gDNA was sheared using the M220 Focused-Ultrasonicater (Covaris, Woburn, MA), followed by end repair, size selection (350-bp insert size), dA-tailing, adaptor ligation, and library amplification by eight rounds of PCR. The quality of the DNA libraries was assessed by Fragment Analyzer with the High Sensitivity NGS Fragment Analysis Kit (Advanced Analytical Technologies, Ankeny, IA). Library concentrations were measured by the StepOnePlus Real-Time PCR System (ThermoFisher Scientific, Waltham, MA) with the KAPA Library Quantification Kit (Kapabiosystems, Wilmington, MA). Each library was labeled with an index sequence during the adaptor-ligation step using Illumina’s TruSeq Dual Indexing system (Illumina, San Diego, CA), which contains 96 unique indices. The 120 libraries were split and pooled into two groups at equal molar concentrations, with each library having a unique index within its group. The library pools were diluted to 2 nM with Reticulocyte Standard Buffer (RSB; 10 mM Tris-HCl, pH 8.5) and denatured with 0.1 M NaOH. Eighteen pM libraries were clustered in a high-output flow cell using the HiSeq Cluster Kit v4 on a cBot (Illumina, San Diego, CA). After cluster generation, the flow cell was loaded onto HiSeq 2500 for sequencing using the HiSeq SBS kit v4 (Illumina, San Diego, CA). DNA was sequenced with paired-end, 100-bp chemistry among four lanes. Raw bcl files were converted to fastq files and demultiplexed using the program bcl2fastq2.17.1.14 (https://github.com/thepler/bcl2fastq2).

#### Mapping and genotyping

Raw reads were trimmed to remove adapters with Trimgalore! *(--quality 0*).^88^ Trimmed reads for all 120 individuals were mapped to the *Xiphophorus maculatus* reference genome (v.5.0, RefSeq accession number: GCF_002775205.1)^89^ using BWA-MEM.^90^ We used this reference genome because *X. maculatus* was the only species in the family Poeciliidae with a chromosome-scale genome assembly. On average, ∼94% of trimmed reads mapped across all individuals (Table S4). After mapping, based on percent of mapped reads, it became clear that one sample (sample MX16-205) labeled as *Pseudoxiphophorus* from a sulfidic habitat was likely mislabeled, and this sample was excluded from all further analyses. Note that we conducted the same analyses mapping to the *P. mexicana* reference genome, and all results were consistent.

The resequenced genomes generated for this study had an average sequence depth of 3.8 (± 0.4; Table S4). We used the Genome Analysis Toolkit (GATK, v.3.5)^100^ and Picard Tools^101^ to mark duplicate reads (using the MarkDuplicates tool) and realign mapped reads around indels (using the RealignerTargetCreator and IndelRealigner tools) following GATK best practices.^102,103^ The resulting realigned bam-file for each individual was used in subsequent analyses. GATK was also used to identify single nucleotide polymorphisms (SNPs) per population. HaplotypeCaller was used to identify genotypes per individual with the *--ERC* GVCF flag.^100^

Samples were genotyped jointly on a per-species basis using GATK’s GenotypeGVCFs tool after combining GVCF files from the sulfidic and nonsulfidic populations using the GATK CombineGVCFs tool. Genotypes were identified using the *--include-non-variant-sites* flag to retain all sites in the initial VCF. We recoded genotypes inferred as phased missing (.|.) as unphased missing (./.) due to a formatting issue with VCFtools. We used the GATK SelectVariants to extract biallelic SNPs from the original genotyped GVCF files, except for the *d*_XY_ analysis (see below). For all analyses relying on biallelic SNPs, combined VCF files with both the sulfidic and nonsulfidic populations of each species were filtered to exclude sites with >25% missing data, a minor allele frequency below 5%, and alleles that were fixed for the non-reference allele were removed using VCFtools v.0.1.17 with the following flags: *--max-missing* 0.75, *--remove-filtered-all*, *--maf 0.05*, and *--max-non-ref-af-any* 0.999. After filtering, 1,994,206 biallelic SNPs were retained for *Poecilia*, 5,284,684 for *Pseudoxiphophorus*, and 4,624,801 for *Xiphophorus*. For calculating *d*_XY_, only indels and sites with >25% missing data were removed from the merged VCF using VCFtools with the *-remove-indels* and *-max-missing 0.75* flags, respectively (note that this filtering retained monomorphic sites for accurate estimation of *d*_XY_).^104^ This resulted in the retention of 439,564,228 monomorphic and biallelic sites for *Poecilia*, 528,674,992 for *Pseudoxiphophorus*, and 611,098,475 for *Xiphophorus*.

#### Analyzing population structure

We assessed structure between populations in sulfidic and nonsulfidic habitats of each species. To do so, we removed sites with an *r*^2^ above 0.7 within 5 kb within VCFtools *(--geno-r2 --min-r2* 0.7 *--ld-window* 5000), producing a set of unlinked SNPs for each species (62,738 in *Poecilia*, 64,405 in *Pseudoxiphophorus*, and 65,515 in *Xiphophorus*). We investigated the extent of gene flow between populations using FastStructure,^105^ a maximum-likelihood-based assignment method that estimates admixture proportions for each individual. FastStructure was run with the number of populations (*K*) set as 2 to test if individuals cluster by habitat of origin (*i.e.*, sulfidic vs. nonsulfidic). Population structure was further examined by principal component analysis (PCA) based on a variance-standardized genetic relationship matrix computed with *PLINK* v.2.00.^106^

From the structure analyses, we identified an admixed individual of *P. mexicana* that was likely an F1 hybrid between the sulfidic and non-sulfidic populations (MX16-023). We also identified another admixed individual in *P. mexicana* that was possibly an F1 or F2 backcross (MX16-121). We removed these admixed individuals from the *Poecilia* SNP sets for downstream analyses.

#### Reciprocal translocation experiments

To experimentally validate local adaptation^32^ and test for natural selection against migrants between sulfidic and nonsulfidic populations,^33^ we conducted reciprocal translocation experiments using previously established approaches.^19,107^ Large (20-liter) plastic buckets were placed into the two habitat types as experimental mesocosms. Two holes (18 × 32 cm) were cut into opposite sides of each bucket and sealed with 1.5 mm plastic mesh to allow the free exchange of water with the environment. Approximately 50 small holes (<1 mm) were drilled into the bucket lids to facilitate air exchange. Mesocosms were placed into shallow areas of the sulfidic and nonsulfidic streams, seeded with a 3–4 cm layer of natural substrate, and allowed to settle overnight before the start of the experiment. We established ten mesocosms in each habitat. Water conditions in these mesocosms have been shown to closely match conditions in the surrounding stream.^107^

We collected adult *Poecilia*, *Pseudoxiphophorus*, and *Xiphophorus* by seine and placed them in insulated coolers with aerated water for transport to the mesocosm locations. Five haphazardly chosen individuals of the same species and habitat type were introduced into a mesocosm. Half of the mesocosms in each habitat were used as controls to test the survival of resident fish, and the other half to test survival of fish from the opposite habitat type. Transportation and handling times were minimal (<1 hour) and were balanced for resident and translocated individuals. Fish were measured for standard length prior to introduction. Experiments ran for ∼20 hours before termination.

Following the experimental period, we quantified survival and returned surviving individuals to their original collection site.

To analyze variation in survival, we used a generalized linear mixed model (GLMM) with a binomial error distribution and a logit-link function, as implemented in the R package LME4.^108^ Survival (binary data: 0 = died; 1 = survived) was used as the response variable. We included species, habitat of origin (sulfidic vs. nonsulfidic), and testing habitat (sulfidic vs. non-sulfidic) as fixed effects (including their interactions terms), and standard length (log_10_-transformed) as a covariate. Additionally, mesocosm ID was included as a random effect.

### Evidence for convergence at a genomic level

The final objective of this study was to identify genomic signatures of convergent evolution. To do so, we leveraged the genome resequencing data described above. We analyzed patterns of molecular evolution in the mitochondrial genomes of focal populations and explored patterns of divergence and signatures of selection in the nuclear genomes as well.

#### Mitochondrial genome assembly and analysis

Trimmed reads from the whole-genome re-sequencing were used as input for NOVOPlasty,^109^ which allows for *de novo* assembly and variant-calling in short circular genomes. The *X. maculatus* reference mitochondrial genome (NCBI Accession NC_011379) was used as the seed input and reference sequence. Sequences for the 13 protein-coding genes were identified using MitoAnnotator.^110^

To investigate the evolutionary history of mitochondrial genomes among populations, we performed phylogenetic analyses using 12 of the 13 protein-coding mitochondrial genes (*atp8* was excluded from analyses due to poor assembly for this short gene in many individuals). For each gene, we aligned nucleotide sequences of all individuals using the default options of the MAFFT progressive alignment method,^111^ guided by a translated amino acid sequence with TranslatorX.^112^ We used PopART v.1.7^113^ to construct a TCS haplotype network^114^ for the concatenated mitochondrial sequences of each population pair separately.

Since mitochondria and OxPhos are the direct target of H_2_S toxicity, we analyzed patterns of molecular evolution and the effect of positive selection in lineages from sulfidic habitats for 12 of the 13 mitochondrially encoded protein-coding genes, following a previously developed approach.^25^ In brief, because of the high coverage of the RNAseq data, we used GATK^100^ to realign BWA-MEM mapped RNAseq reads around indels and used the UnifiedGenotyper to detect variants following GATK best practices.^102,103^ We genotyped individuals on a per-population basis using the UnifiedGenotyper EMIT_ALL_SITES output mode to create a VCF-file for each population. We then used a custom pipeline to fill positions lacking sequence data in each VCF with Ns, rather than the GATK default (which fills in missing data with the reference allele).^115^ We used BCFtools v.1.3 to designate the genotype at all variable sites as the major allele in each population’s VCF-file with the setGT tool.^116^ Using the coordinates for each mitochondrial gene from the *X. maculatus* reference genome gene annotation file (GTF), we created population-specific consensus sequences for each mitochondrial gene across all individuals in a population using the consensus tool in BCFtools. We aligned nucleotide sequences for all populations using the MAFFT progressive alignment method^111^ guided by the translated amino acid sequence as implemented in TranslatorX,^112^ producing an in-frame multispecies alignment of each gene. We constructed gene-specific phylogenies (gene trees) for each sequence using the DNAML program in PHYLIP v.3.696.^117^

To identify mitochondrial genes that have experienced positive selection in multiple sulfide spring populations, as evidenced by an excess rate of non-synonymous to synonymous substitutions (ω) for codons in each mitochondrial gene across the sulfidic branches of our phylogenies, we used branch models implemented by codeml in PAML v.4.9a.^118^ For each mitochondrial gene, codeml was run using the in-frame multispecies alignment and gene tree with the three sulfidic populations designated as foreground branches and nonsulfidic populations as background branches, allowing us to test whether ω differed between foreground and background branches in a two-ratio branch model (M2; model = 2, NSsites = 0) compared against a null model with a single ω value for all branches (M0; model = 0, NSsites = 0). A likelihood-ratio test was used to assess whether models significantly differed for each gene, with the test statistic calculated as two times the difference between the ln-likelihood of the branch model and the ln-likelihood of the null model, and *P*-values calculated from a c^2^ approximation.^119^

#### Characterizing the genomic landscape of divergence

To characterize the landscape of genomic divergence, we calculated genomic differentiation (*F*_ST_, relative divergence) and divergence (*d*_XY_, absolute divergence) between the sulfidic and nonsulfidic population of each species across non-overlapping 5-kb windows spanning the genome from our filtered dataset (excluding the two putative, early-stage hybrids in *Poecilia*, as described above) using the popgenWindows.py script.^120–122^ Divergent selection between habitats should lead to elevated levels of both *F*_ST_ and *d*_XY_ at and around loci under selection (those contributing to the maintenance of adaptive divergence), while regions of the genome with reduced diversity in one or both populations (likely a consequence of background selection or low recombination rates) are expected to exhibit elevated *F*_ST_ but not *d*_XY_.^34,37,123^ To identify probable targets of divergent selection between sulfidic and nonsulfidic populations, we used a composite outlier analysis of both statistics as implemented in the R package MINOTAUR.^38^ First, we converted *F*_ST_ and *d*_XY_ to rank-based *P*-values using a right-tailed test, then –log_10_-transformed these *P*-values, and calculated the Mahalanobis distances among windows (the “Md-rank-P” method recommended for summary statistics).^39^ We considered windows in the top 1% of the resulting Mahalanobis distance distribution as genomic regions likely experiencing divergent natural selection between habitat types in each species. We also identified regions that were classified as outlier windows across all three species and between each pairwise combination of species.

We investigated the link between divergent selection and the putative biological functions of genes found in these outlier windows for each species using a GO-enrichment analysis (as described above). We first identified genes in outlier regions from the *X. maculatus* genome annotation file (GTF) using the *intersect* tool in BEDTools v.2.25.0.^124^ We then annotated all genes that had a match in the human SWISSPROT database (21,891 genes) with GO IDs and tested for enrichment of specific GO IDs in outlier regions relative to all human annotated genes in the *X. maculatus* genome using GORILLA (*P*-value threshold: 0.0001, accessed 09/29/2020).^99^ In total, 12,247 unique SWISSPROT annotations were associated with a GO term.

#### Scans for positive selection

To further test for signals of natural selection driving genomic divergence between sulfidic and nonsulfidic populations, we implemented the cross-population composite likelihood ratio (XP-CLR)^40^ test to identify regions of the genome with evidence for selective sweeps in the sulfide spring population of each species based on differentiation in multilocus allele frequencies compared to the nonsulfidic population. We further filtered our biallelic SNP data to include only sites with no more than 10% missing data (957,379 in *Poecilia*, 2,885,156 in *Pseudoxiphophorus*, and 3,419,489 in *Xiphophorus*), and limited our analyzes to chromosomes in the assembly, excluding all short scaffolds. For XP-CLR, the local two-dimensional site-frequency-spectrum (SFS) was compared to the genome-wide two-dimensional SFS to identify genomic regions putatively under selection.^40^ For each species, we designated the nonsulfidic population as the reference population, and the sulfidic population was the target population for each of the comparisons. Prior to running XP-CLR, we partitioned the genome into discrete 25-kb windows. If a window contained more than 200 SNPs, only the first 200 SNPs were used to calculate the composite likelihood ratio. The XP-CLR test statistic was not calculated if a window contained fewer than ten SNPs. We considered windows within the top 1% as putative selective sweeps.

## Data Availability

All sequence data are available at the National Center for Biotechnology Information (NCBI) (https://www.ncbi.nlm.nih.gov under BioProject accession no. PRJNA738871).

## Acknowledgements

We are indebted to the community of Teapa, especially the owners of Rancho La Gloria, for providing access to study sites, and to the Centro de Investigación e Innovación para la Enseñanza y Aprendizaje (CIIEA) for their hospitality and support during the nearly decade-long course of this project. We thank N. Barts, C. Carson, Z. Culumber, T. Doumas, G. W. Hopper, C. N. Passow, and E. A. Renner for assistance in the field, as well as O. Cornejo, K. B. Gido, A. G. Hope, D. A. Marques, and T. J. Morgan for their comments and discussions. This work was supported by grants from the NSF (IOS-1557860, IOS-1931657, and IOS-2311366) and the US Army Research Office (W911NF-15-1-0175, W911NF-16-1-0225) to JLK and MT. RG was supported by the American Museum of Natural History (Theodore Roosevelt Memorial Fund grant), Friends of Sunset Zoo (Conservation Scholar Program), American Livebearer Association (Vern Parish Fund award), Society for the Study of Evolution (Rosemary Grant Advanced Graduate Research Excellence Grant), American Society of Naturalists (Student Research Award), and a NSF Graduate Research Fellowship (DGE-1746899). MT was supported by the Des Lee Collaborative Vision in Zoological Studies.

