## Supplementary material for "Integrative analyses of convergent adaptation in sympatric extremophile fishes"

### Supplementary Tables and Figures

**Table S1.** Results of Procrustes analysis of variance examining variation in body shape in *Poecilia*, *Pseudoxiphophorus*, and *Xiphophorus* from sulfidic and nonsulfidic populations.

df = degrees of freedom
SS = Procrustes distance sum of squares
MS = mean sum of squares
F = *F*-ratio values
Z = effect sizes
P = P-value

| | df | SS | MS | $R^2$ | *F* | *Z* | *P* |
|---|---|---|---|---|---|---|---|
| Centroid size | 1 | 0.020 | 0.020 | 0.004 | 7.939 | 4.782 | <0.001 |
| Species | 2 | 0.054 | 0.027 | 0.011 | 10.907 | 7.308 | <0.001 |
| Habitat | 1 | 0.006 | 0.006 | 0.001 | 2.286 | 2.093 | 0.018 |
| Sex | 1 | 0.048 | 0.048 | 0.010 | 19.260 | 6.362 | <0.001 |
| Centroid size × species | 2 | 0.015 | 0.007 | 0.003 | 2.994 | 3.437 | <0.001 |
| Centroid size × habitat | 1 | 0.002 | 0.002 | 0.000 | 0.929 | 0.066 | 0.473 |
| Centroid size × sex | 1 | 0.006 | 0.006 | 0.001 | 2.500 | 2.257 | 0.012 |
| Species × habitat | 2 | 0.069 | 0.034 | 0.014 | 13.780 | 7.565 | <0.001 |
| Species × sex | 2 | 0.113 | 0.057 | 0.023 | 22.692 | 7.994 | <0.001 |
| Habitat × sex | 1 | 0.005 | 0.005 | 0.001 | 2.013 | 1.811 | 0.035 |
| Species × habitat × sex | 2 | 0.015 | 0.007 | 0.003 | 2.970 | 3.531 | <0.001 |
| Residuals | 407 | 1.015 | 0.002 | 0.210 | | | |
| Total | 423 | 4.833 | | | | | |

**Table S2.** List of up- and downregulated genes in *Poecilia*, *Pseudoxiphophorus,* and *Xiphophorus* from the sulfidic habitat. The final sheet indicates the shared differentially expressed genes. Blue rows are upregulated genes, and red are downregulated genes. This table is provided in a separate file: "Table S2 – Differential Expression.xlsx". The table includes the gene number from the *X. manulatus* reference annotation (Gene Number X. maculatus), the corresponding *X. maculatus* geneID (X. maculatus GeneID), the human gene ID if one exists (HumanGene), the BLAST e-value for identifying the closest human homolog (BLAST E value), the $log_2$ fold change between sulfidic and nonsulfidic samples (logFC), log counts per million (logCPM), likelihood ratio (LR), p-value (P Value) and false discovery rate calculated with the Benjamini-Hochberg correction (FDR).

**Table S3.** Results of the GO enrichment analysis of differentially expressed genes in *Poecilia, Pseudoxiphophorus, and Xiphophorus*, including a list of GO terms with evidence for significant enrichment (the identification number and description of each GO term), the total number of genes in the reference set (N), the total number of genes associated with a specific GO term (B), the number of genes in the target set (upregulated or downregulated genes; n), and the number of genes in the intersection (b). Enrichment was calculated from these values ([b/n]/[B/N]). Also included is the P-value associated with enrichment and the false discovery rate adjusted significance (q-value). The final sheet indicates shared enriched GO terms. This table is provided in a separate file: "Table S3 – GO Enrichment.xlsx".

**Table S4.** Average read counts after quality control, average percentage of quality-controlled reads that mapped, and the average depth of coverage of genomic DNA reads mapped to the *X. maculatus* reference genome per population (*N* = 19–20 per population).

| Species | Habitat | Paired Reads After Trimming | % Paired Reads Aligned | Depth of Coverage |
|---|---|---|---|---|
| *P. mexicana* | NS | 32,251,543.60 | 87.5 | 3.33 |
| *P. mexicana* | S | 30,998,540.60 | 87.6 | 3.20 |
| *P. bimaculatus* | NS | 30,295,782.70 | 95.5 | 3.75 |
| *P. bimaculatus* | S | 32,404,180.11 | 95.6 | 4.02 |
| *X. hellerii* | NS | 31,606,354.70 | 98.5 | 4.32 |
| *X. hellerii* | S | 30,549,363.70 | 98.6 | 4.19 |

**Table S5.** Results from translocation analysis. Data were analyzed with a Binomial Generalized Linear Mixed Model with mesocosm ID as a random effect.

SE = standard error
*Z* = z-score
*P* = *P*-value

| Fixed effects | Estimate | SE | *z* | *P* |
|---|---|---|---|---|
| (Intercept) | -3.986 | 4.090 | -0.975 | 0.330 |
| $\log_{10}$(standard length) | 4.888 | 2.664 | 1.835 | 0.067 |
| **Habitat of origin** | **-2.368** | **1.015** | **-2.333** | **0.020** |
| **Test habitat** | **-6.722** | **1.128** | **-5.962** | **<0.001** |
| Species (*X. hellerii*) | 1.233 | 1.363 | 0.904 | 0.366 |
| Species (*P. mexicana*) | -1.538 | 1.115 | -1.379 | 0.168 |
| **Habitat of origin × Test habitat** | **7.470** | **1.082** | **6.905** | **<0.001** |
| Habitat of origin × Species (*X. hellerii*) | -0.578 | 1.360 | -0.425 | 0.671 |
| Habitat of origin × Species (*P. mexicana*) | 1.580 | 1.217 | 1.299 | 0.194 |
| Test habitat × Species (*X. hellerii*) | 0.762 | 1.171 | 0.650 | 0.516 |
| Test habitat × Species (*P. mexicana*) | 0.917 | 1.134 | 0.808 | 0.419 |

**Table S6.** Results of analyses of molecular evolution in mitochondrial genes using branch tests to evaluate differences in non-synonymous amino acid substitution rates (ω) between nonsulfidic (background; $\omega_{background}$) and sulfidic (foreground, $\omega_{foreground}$) populations. We provide ω values of the null model ($\omega_{nonsulfidic} = \omega_{sulfidic}$; $\omega_{null}$) when a two ω branch model was not supported. Genes with evidence for positive selection ($\omega_{nonsulfidic} \neq \omega_{sulfidic}$) are indicated in bold. Log likelihoods of the null (lnLnull) and test (lnLtest) statistics, 2*change in loglikelihood (2ΔlnL) and P-value (*P*) are also reported.

| Gene | $lnL_{null}$ | $lnL_{test}$ | 2ΔlnL | *P* | $\omega_{null}$ | $\omega_{background}$ | $\omega_{foreground}$ |
|---|---|---|---|---|---|---|---|
| ATP6 | -418.20 | -418.16 | 0.08 | 0.780 | 0.019 | | |
| ATP8 | -186.87 | -186.87 | 0.00 | 1.000 | 1.000 | | |
| **COX1** | **-3337.63** | **-3316.92** | **41.43** | **<0.001** | | **0.020** | **999.000** |
| COX2 | -1317.27 | -1317.57 | 0.61 | 0.435 | 0.018 | | |
| **COX3** | **-1550.41** | **-1537.91** | **25.00** | **<0.001** | | **0.021** | **999.000** |
| **CYTB** | **-2096.27** | **-2088.36** | **15.82** | **<0.001** | | **0.025** | **999.000** |
| ND1 | -1576.02 | -1574.27 | 3.49 | 0.062 | 0.017 | | |
| **ND2** | **-1474.38** | **-1470.64** | **7.47** | **0.006** | | **0.040** | **0.203** |
| **ND3** | **-738.04** | **-733.09** | **9.91** | **0.002** | | **0.030** | **999.000** |
| ND4 | -2113.73 | -2113.35 | 0.78 | 0.379 | 0.030 | | |
| ND4L | -634.96 | -633.93 | 2.05 | 0.152 | 0.032 | | |
| ND5 | -2699.69 | -2699.15 | 1.09 | 0.296 | 0.021 | | |
| ND6 | -558.09 | -558.26 | 0.35 | 0.555 | 0.012 | | |

**Table S7.** List of genes found in regions of elevated genomic divergence (Mahalanobis distance of $F_{ST}$ and $d_{XY}$) between sulfidic and nonsulfidic populations of *Poecilia*, *Pseudoxiphophorus*, and *Xiphophorus*. Provided are the locations of each outlier window and each gene in the *X. maculatus* reference genome, the NCBI Gene ID of each gene in the *X. maculatus* reference genome, as well as the accession number and gene name based on a BLAST search against the SwissProt database of human genes. Outlier genes found in each species are denoted by an x in the relevant species column. This table is provided in a separate file: "Table S7 - Md outliers.xlsx".

**Table S8.** Results of the GO enrichment analysis of genes found in outlier regions of genomic divergence (Mahalanobis distance of $F_{ST}$ and $d_{XY}$) between sulfidic and nonsulfidic *Poecilia*, sulfidic and nonsulfidic *Pseudoxiphophorus* , sulfidic and nonsulfidic *Xiphophorus* , sulfidic and nonsulfidic *Poecilia* and *Pseudoxiphophorus*, sulfidic and nonsulfidic *Poecilia* and X*iphophorus, and* sulfidic and nonsulfidic *Pseudoxiphophorus* and *Xiphophorus,* including a list of GO terms with evidence for significant enrichment. Parameters provided are the same as described in Table S3. This table is provided in a separate file: "Table S8 - Outlier GO enrichment.xlsx".

**Table S9.** List of genes found in regions with evidence of a selective sweep (XP-CLR) between sulfidic and nonsulfidic populations of *Poecilia*, *Pseudoxiphophorus*, and *Xiphophorus*. Provided are the locations of each outlier window and each gene in the *X. maculatus* reference genome, the NCBI Gene ID of each gene in the *X. maculatus* reference genome, as well as the accession number and gene name based on a BLAST search against the SwissProt database of human genes. Outlier genes found in each species are denoted by an x in the relevant species column. This table is provided in a separate file: "Table S9 - XP-CLR sweeps.xlsx".

**Table S10.** Results of the GO enrichment analysis of genes found in genomic regions found to have experienced selective sweeps in sulfidic *Pseudoxiphophorus* (XP-CLR) and results of the GO enrichment analysis of genes found in genomic regions found to have experienced selective sweeps in sulfidic *Xiphophorus* (XP-CLR), including a list of GO terms with evidence for significant enrichment. Parameters provided are the same as described in Table S3. This table is provided in a separate file: "Table S10 - Pbimac XP-CLR Xhell XP-CLR GO enrichment.xlsx".

**Table S11.** List of collection locations and final sample sizes for different aspects of this study.

| Locality | $H_2S$ | $N_{genomes}$ | $N_{translocation}$ (resident/migrant) | $N_{morphology}$ | $N_{tolerance}$ | $N_{RNAseq}$ |
|---|---|---|---|---|---|---|
| La Gloria springs, Rio Pichucalco drainage, Chiapas, Mexico (17.532, -93.015) | + | | | | | |
| *Poecilia mexicana* | | 20 | 25/25 | 52 | 12 | 6 |
| *Pseudoxiphophorus bimaculatus* | | 19 | 25/25 | 63 | 12 | 6 |
| *Xiphophorus hellerii* | | 20 | 25/25 | 99 | 12 | 6 |
| Arroyo Caracol, Rio Pichucalco drainage, Chiapas, Mexico (17.537, -93.017) | - | | | | | |
| *Poecilia mexicana* | | 20 | 25/25 | 115 | 16 | 6 |
| *Pseudoxiphophorus bimaculatus* | | 20 | 25/25 | 52 | 14 | |
| *Xiphophorus hellerii* | | 20 | 25/24 | 43 | 12 | 6 |
| Arroyo Pujil, Rio Ixtapangajoya drainage, Chiapas, Mexico (17.476, -92.986) | - | | | | | |
| *Pseudoxiphophorus bimaculatus* | | | | | | 6 |

**Table S12.** Average read counts after quality control and the average percentage of quality-controlled reads that mapped to the *X. maculatus* reference genome for each of the population ($N = 6$ per population).

| Species | Habitat | Paired Reads After Trimming | % Paired Reads Aligned |
|---|---|---|---|
| *P. mexicana* | NS | 83,042,097 | 97.0 |
| *P. mexicana* | S | 76,339,057 | 96.3 |
| *P. bimaculatus* | NS | 35,247,808 | 96.9 |
| *P. bimaculatus* | S | 39,545,078 | 95.7 |
| *X. hellerii* | NS | 30,761,274 | 98.8 |
| *X. hellerii* | S | 29,296,035 | 99.0 |

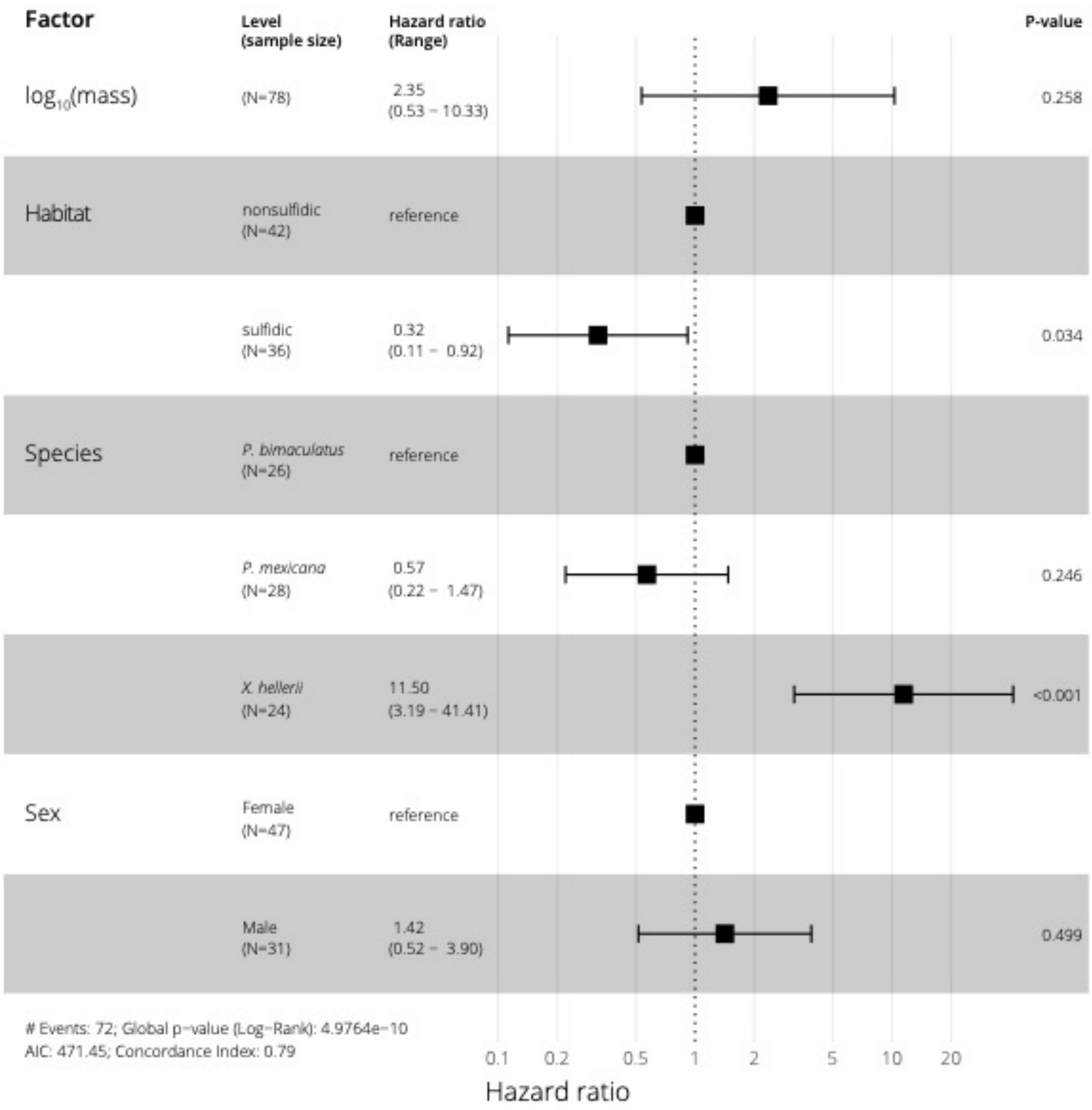


**Figure S1.** Forest plot of Cox regression for time to loss of equilibrium upon acute exposure to $H_2S$ as a measure of sulfide tolerance. Body mass ($log_{10}$-transformed), habitat of origin, species, and sex served as predictor variables.

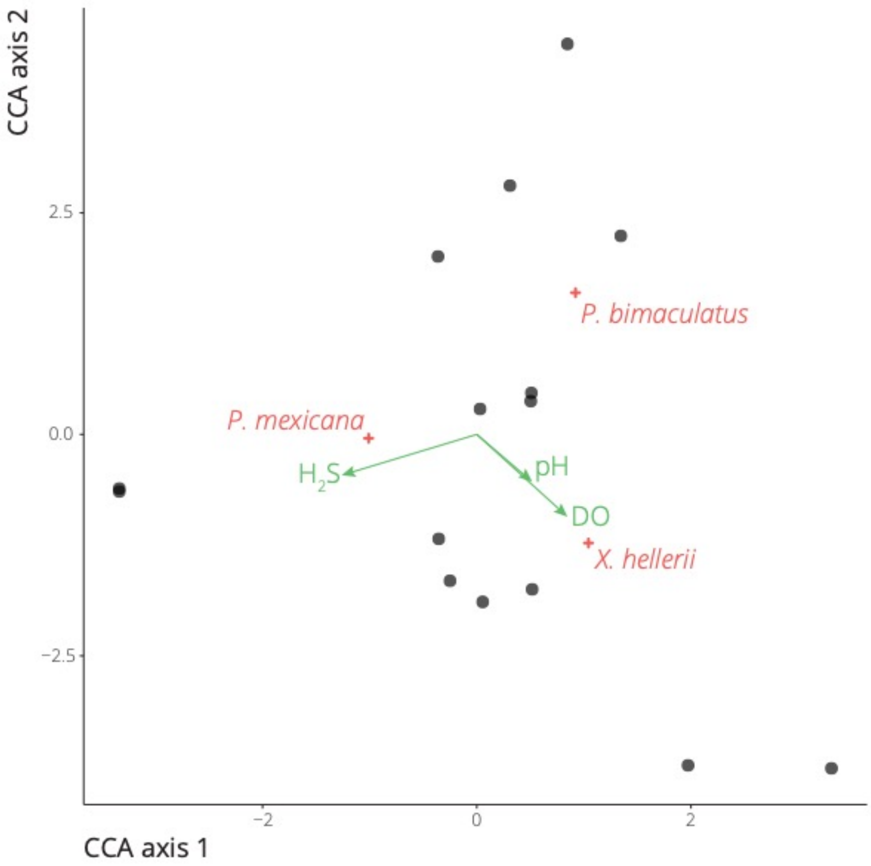


**Figure S2.** Results of a canonical correspondence analysis (CCA) that investigated the relative abundance of the focal species across 15 sites within the La Gloria sulfide spring complex as a function of water chemistry. Community composition was estimated by seining stream segments and counting members of each species. Water parameters (temperature, specific conductivity, pH, dissolved oxygen, and $H_2S$ concentration) were quantified at each site. The variance inflation factors (VIF) of variables were checked with the USDM package in R, variables with VIF > 5 were excluded from the final CCA. The final CCA model was significant ($F_{3,13} = 3.01$, $P = 0.047$), with significant effects on relative abundance by dissolved oxygen ($F_{1,13} = 4.13$, $P = 0.043$) and $H_2S$ concentrations ($F_{1,13} = 4.89$, $P = 0.023$), but not pH ($F_{3,13} < 0.01$, $P = 0.998$).

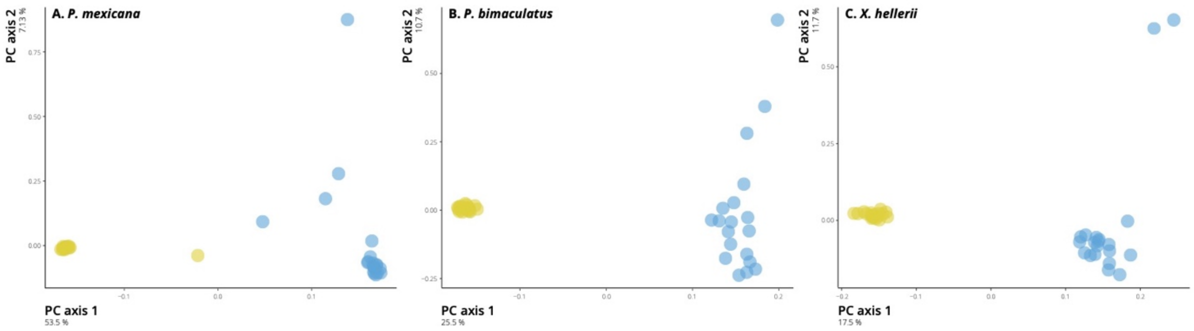


**Figure S3.** Principal component analyses of genomic variation among all sampled individuals from the sulfidic (yellow) and the nonsulfidic (blue) site for (A) *Poecilia*, (B) *Pseudoxiphophorus*, and (C) *Xiphophorus* based on unlinked SNPs. The percentage of variance explained by each principal component is indicated on the axis.

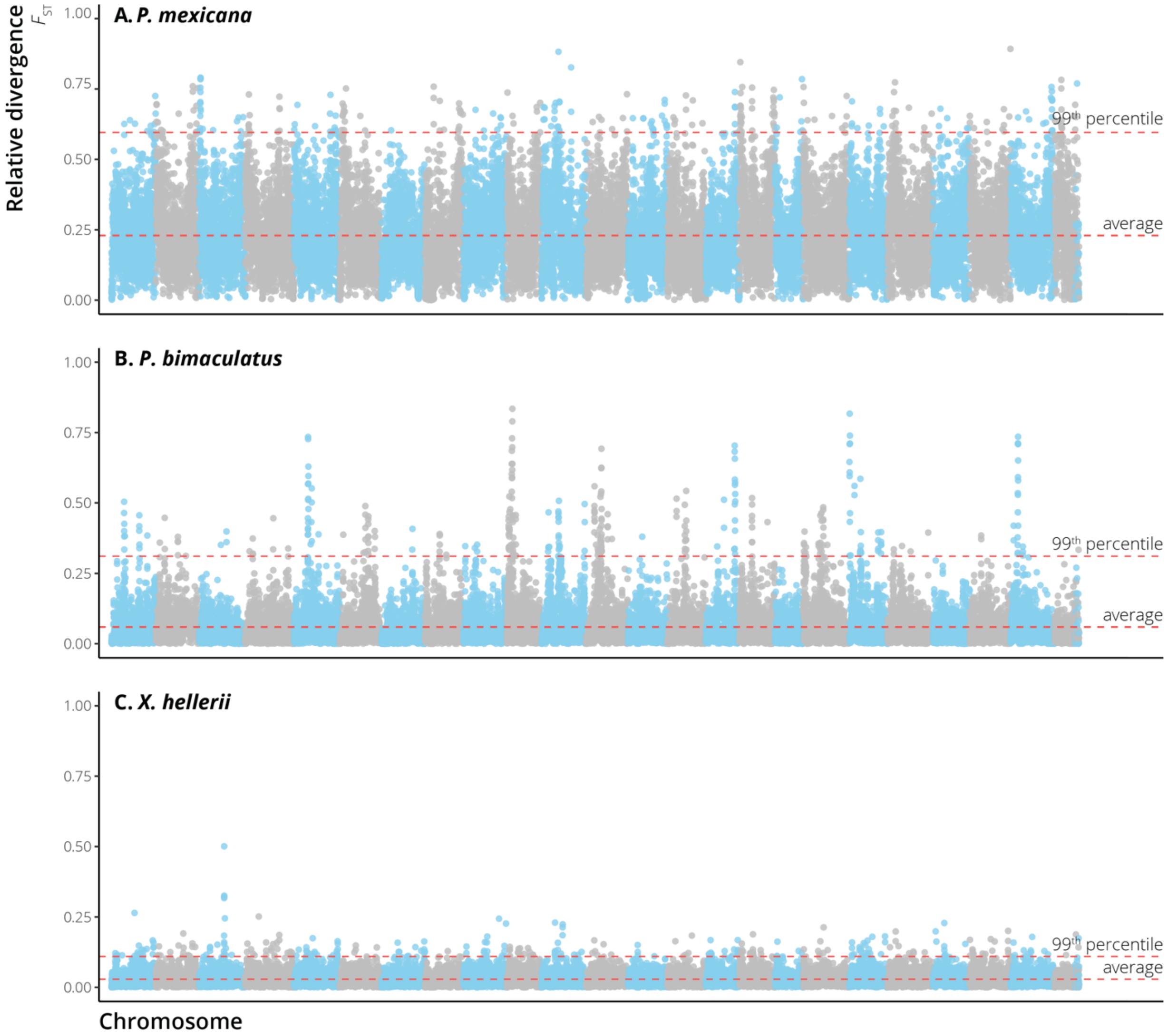


**Figure S4.** Relative genomic divergence ($F_{ST}$) between the nonsulfidic and sulfidic population of each species measured for non-overlapping 25-kb windows across the genome. The 99th percentile and average $F_{ST}$ are indicated by red dashed lines.

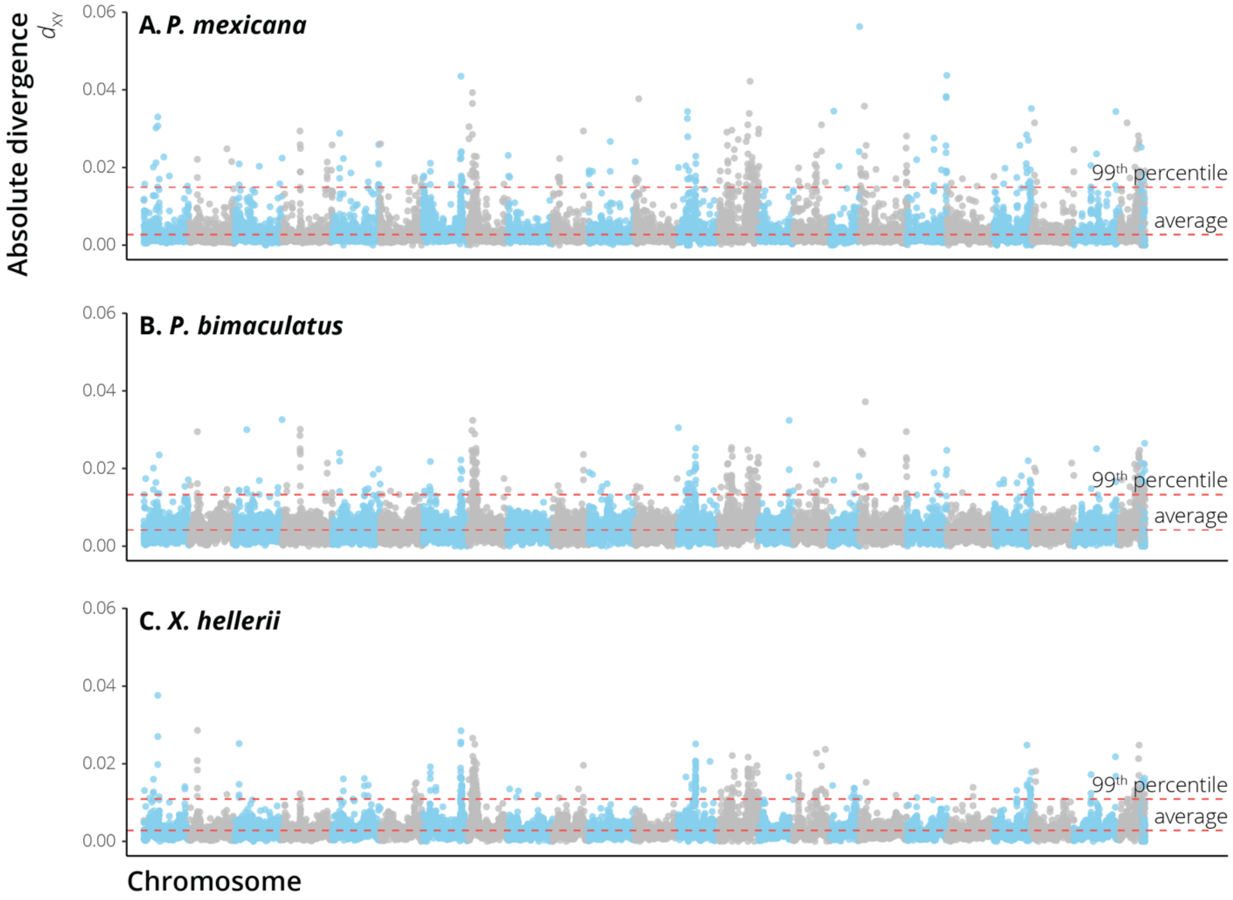


**Figure S5.** Absolute genomic divergence ($d_{XY}$) between the nonsulfidic and sulfidic population of each species measured for non-overlapping 25-kb windows across the genome. The 99$^{th}$ percentile and average $d_{XY}$ are indicated by red dashed lines.

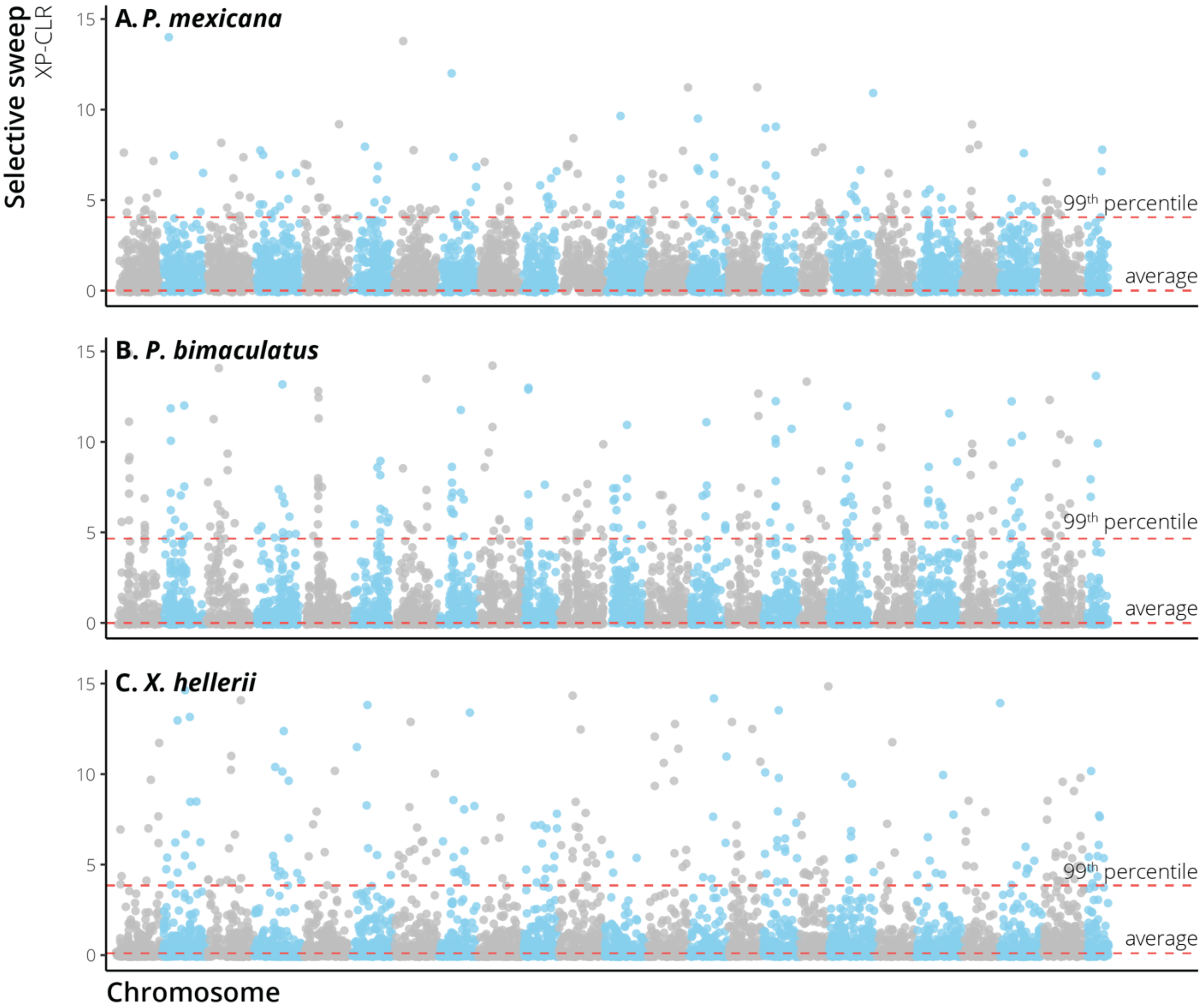


**Figure S6.** Allele frequency spectrum-based estimates of selective sweeps (XP-CLR) in the sulfidic population across the genome. The 99th percentile and average XP-CLR score are indicated by red dashed lines.

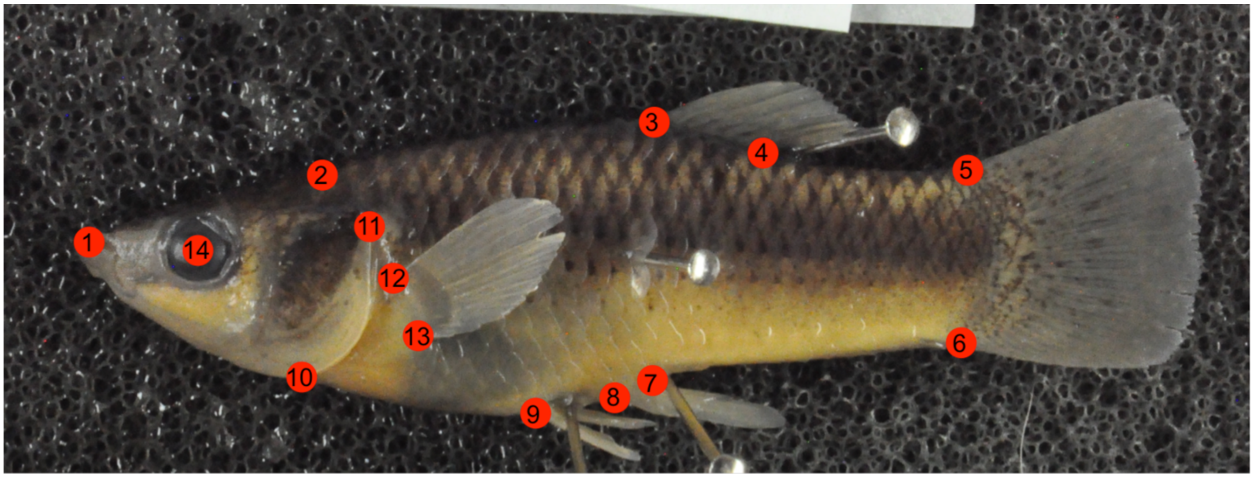

**Figure S7.** Landmark locations used for geometric morphometric analyses. Landmarks included the tip of the upper jaw (1); the posterior head region (2); the anterior (3) and posterior (4) insertions of the dorsal fin; the dorsal (5) and ventral (6) insertions of the caudal fin; the posterior (7) and anterior (8) insertions of the anal fin; the anterior insertion of the pelvic fin (9); the bottom of the head where the operculum breaks away from the body outline (10); the posterodorsal corner of the operculum (11); the ventral (12) and dorsal (13) insertion of the pectoral fin; and the center of the eye (14).
